# Identification of QTL hotspots affecting agronomic traits and high-throughput vegetation indices in rainfed wheat

**DOI:** 10.1101/2021.06.25.449881

**Authors:** Rubén Rufo, Andrea López, Marta S. Lopes, Joaquim Bellvert, Jose Miguel Soriano

## Abstract

Understanding the genetic basis of agronomic traits is essential for wheat breeding programmes to develop new cultivars with enhanced grain yield under climate change conditions. The use of high-throughput phenotyping (HTP) technologies for the assessment of agronomic performance through drought-adaptive traits opens new possibilities in plant breeding. HTP together with a genome-wide association study (GWAS) mapping approach can become a useful method to dissect the genetic control of complex traits in wheat to enhance grain yield under drought stress. This study aimed to identify molecular markers associated with agronomic and remotely sensed vegetation index (VI)-related traits under rainfed conditions in bread wheat and to use an *in silico* candidate gene (CG) approach to search for upregulated CGs under abiotic stress. The plant material consisted of 170 landraces and 184 modern cultivars from the Mediterranean basin that were phenotyped for agronomic and VI traits derived from multispectral images over three and two years, respectively. GWAS identified 2579 marker–trait associations (MTAs). The QTL overview index statistic detected 11 QTL hotspots involving more than one trait in at least two years. A candidate gene analysis detected 12 CGs upregulated under abiotic stress in 6 QTL hotspots. The current study highlights the utility of VI to identify chromosome regions that contribute to yield and drought tolerance under rainfed Mediterranean conditions.

## Introduction

Wheat (*Triticum aestivum* L.) is the most common cultivated crop worldwide. It is grown on 216 million hectares of land, and its global production of 765 million tons of grain provides 19% of the calories and 21% of the protein in the human diet (Faostat 2019, http://www.fao.org/faostat). To cover the expected food demand of a world population that will increase up to 60% by 2050, wheat production needs to increase by 1.7% per year (Leegood et al. 2010). Achieving this objective will not be easy considering the expected negative effects of climate change on wheat yield, particularly in areas such as the Mediterranean basin, where a rise in temperatures by 3–5°C and a decrease in annual rainfall by 25–30% have been predicted (Giorgi and Lionello 2008). An increasing frequency and severity of terminal drought stress will reduce grain weight, grain quality, and wheat yield (Araus et al. 2002; Slafer et al. 2005; Kulkarni et al. 2017). Therefore, there is a need to improve the identification of genotypes able to maintain acceptable levels of yield and yield stability in semiarid environments, which have been identified as the most sensitive to the effects of climate change (Rufo et al. 2021). The release of improved cultivars with enhanced drought adaptation will be critical for breeding programmes focusing on wheat adaptability and stability under rainfed conditions (Graziani et al. 2014; Bhatta et al. 2018).

The recent progress in high-throughput phenotyping (HTP) based on the use of multispectral images acquired from unmanned aerial vehicles (UAVs) has increasingly improved the assessment of agronomic traits (Gracia-Romero et al. 2017; Xie and Yang 2020; Gomez-Candon et al. 2021; Rufo et al. 2021) on large germplasm collections in a rapid, cost-effective, and high spatial resolution way (Duan et al. 2017), as it allows the estimation of various plant traits using nonintrusive and nondestructive technology (White et al. 2012; Rufo et al. 2021). Remote sensing has attracted growing interest in breeding programmes since it can deliver detailed information about biophysical crop traits in many situations to cope with the current phenotyping bottleneck (Araus and Cairns 2014; Juliana et al. 2019; Bellvert et al. 2021). Some studies have demonstrated the use of vegetation indices (VI) to indirectly detect wheat plants under water stress due to a decrease in vegetative growth (Condorelli et al. 2018). Others have demonstrated the use of energy balance models to estimate the actual water status (Gomez-Candon et al. 2021). When VIs are derived from multispectral cameras, they are obtained from the combination of wavelengths located at the visible, red-edge and near-infrared (NIR) regions of the light spectrum (Kyratzis et al. 2017). These wavelengths allow discerning differences in vegetative greenness, rate of senescence, photosynthetic efficiency and stay green duration (Stenberg et al. 2004; Babar et al. 2006; Lopes and Reynolds 2012). It has been stated that anthesis and milk grain are the most suitable growth stages for the assessment of agronomic traits on a plot-by-plot basis (Aparicio et al. 2002; Royo et al. 2003). The use of HTP as a suitable and accurate predictor of agronomic traits such as phenology, grain filling duration, biomass and yield will provide unique opportunities to increase the power of QTL discovery by increasing the number of genotypes included in the analysis (Juliana et al. 2019). This method will increase the frequency of rare alleles of potential interest to improve wheat adaptation to various environmental conditions.

The dissection of the genetic and molecular basis of complex traits such as yield and drought stress tolerance through complementary approaches such as QTL mapping and genome-wide association studies (GWAS) or association mapping is essential in breeding programmes. GWAS is based on linkage disequilibrium (LD) (Flint-Garcia et al. 2003), and it is a powerful approach that provides high mapping resolution due to the higher recombination events analysed in comparison with biparental mapping (Soriano et al. 2017; Qaseem et al. 2019). AM has been used to identify genomic regions related to drought and heat tolerance in durum and bread wheat (Maccaferri et al. 2016; Valluru et al. 2017). Several studies have been conducted to investigate the genetic basis of grain yield and yield-related traits in bread wheat under rainfed conditions using association mapping (Edae et al. 2014; Gizaw et al. 2018; Qaseem et al. 2019; Mérida-García et al. 2020). The release of genome sequences for emmer wheat (Avni et al. 2017), bread wheat (IWGSC 2018) and durum wheat (Maccaferri et al. 2019) and the availability of open databases of RNA-seq experiments (Ramírez-González et al. 2018) have made it possible to use a candidate gene (CG) approach to find targets within QTL intervals without performing new functional studies.

The aim of the current study was to identify molecular markers linked to important agronomic traits, VIs and plant features related to drought resistance assessed by HTP, to define the most important QTL hotspots for such traits and to perform *in silico* detection of the underlying CG in those genomic regions.

## Materials and methods

### Plant material and field trials

A germplasm collection of 354 bread wheat (*Triticum aestivum* L.) genotypes from the MED6WHEAT IRTA panel described in Rufo et al. (2019) was used in this study, of which 170 corresponded to landraces and 184 to modern varieties collected and adapted to 24 and 19 Mediterranean countries, respectively (Online resource 1). The panel is structured into 6 genetic subpopulations (SP) and 38 genotypes that remained admixed (Rufo et al. 2019). SP1: west Mediterranean landraces (43 accessions); SP2: north Mediterranean landraces (59 accessions); SP3: east Mediterranean landraces (42 accessions); SP4: France-Italy modern germplasm (82 accessions); SP5: Balkan modern varieties (24 accessions); and SP6: CIMMYT-ICARDA derived varieties (62 accessions).

The field trials were conducted at Gimenells, Lleida (41°38’ N and 0°22’ E, 260 m.a.s.l), northeastern Spain, under rainfed conditions for three consecutive seasons: 2016, 2017 and 2018. Average minimum and maximum monthly temperatures and rainfall were calculated from daily data recorded for a weather station close to the experimental fields. Experiments followed a non-replicated augmented design with two replicated checks (*cv*. ‘Anza’ and ‘Soissons’) at a ratio of 1:4 between checks and tested genotypes and in 3.6 m^2^ plots with eight rows spaced 0.15 m apart. The sowing density was adjusted to 250 germinable seeds m^-2^, and the sowing dates were 02 December 2015, 21 November 2016, and 15 November 2017. Weeds and diseases were controlled following standard practices at the site.

### Agronomic data

The following traits were measured across the three years according to the protocol described in Rufo et al. (2021): grain yield (GY, t ha^-1^), number of spikes per m^2^ (NSm^2^), number of grains per m^2^ (NGm^2^), thousand kernel weight (TKW, g), aboveground biomass at physiological maturity (biomass, t DM ha^-1^), harvest index (HI), plant height (PH, cm) and early vigour (estimated as green area, GA). The HI was calculated as the ratio between grain and plant weights in a 1-m long row sample. PH was measured at maturity for three main stems per plot and was measured from the soil to the top of the spike, excluding the awns. Early vigour was calculated by integrating the green area (GA) values obtained by ground-based RGB images taken every fourteen days as described in (Casadesús and Villegas 2014) from emergence until the detection of the first node. Finally, days from sowing to anthesis (DSA, GS65) and grain filling duration (GFD, GS87) were measured on each plot based on the growth stage (GS) scale of Zadoks et al. (1974). Growth stages were achieved when at least 50% of the plants in each plot reached them.

### Image acquisition

Image acquisition was conducted with a multispectral camera (Parrot Sequoia) (Parrot, Paris, France) installed onboard an UAV (DJI S800 EVO hexacopter, Nanshan, CHN). Images were acquired on 21 April and 19 May 2017 and 17 April and 18 May 2018. Flights were always conducted at approximately 12:00 solar time under sunny conditions and with a wind speed below 12 m/s. The UAV flew at a height of 40 m agl (above ground level) and with a flight plan of 80/60 frontal and side overlap. The multispectral camera has four spectral bands located at wavelengths of 550 ± 40 nm (green), 660 ± 40 nm (red), 735 ± 10 nm (red edge), and 790 ± 40 (near infrared). The camera yields a resolution of 1280 x 960 pixels. All images were radiometrically corrected through an external incident light sensor that measured the irradiance levels of light at the same bands as the sensor, as well as with in situ spectral measurements in ground calibration targets (black, white, soil and grass). Spectral measurements were conducted with a Jaz spectrometer (Ocean Optics, Inc., Dunedin, FL, USA). Jaz has a wavelength response from 200 to 1100 nm and an optical resolution of 0.3 to 10.0 nm. The calibration of the spectrometer measurements was taken using a reference panel (white colour Spectralon^™^). Geometrical correction was conducted by using ground control points (GCPs) and measuring the position in each with a handheld global positioning system (GPS) (Geo7x, Trimble GeoExplorer series, Sunnyvale, CA, USA). All images were mosaicked using Agisoft Photoscan Professional version 1.6.2 (Agisoft LLC., St. Petersburg, Russia) software and geometrically and radiometrically corrected with QGIS 3.2.0 (USA, http://www.qgis.org). Then, six spectral vegetation indices (VIs) were carefully selected based on their significance in relation to certain plant physiology features in wheat (Table 1). In addition, the leaf area index (LAI) was measured using a portable ceptometer (AccuPAR model LP-80, decagon devices Inc., Pullman, WA, USA) from 13:00 to 15:00 (local time) on each image acquisition date in 64 different plots of each set of landrace and modern set. Then, the LAI was estimated for each plot in the whole collection through the MTVI2, following the methodology described by Rufo et al. (2021) and Gomez-Candón et al. (2021). All VIs were assessed in 2017 and 2018 through UAV multispectral images at two growth stages: 1) when all the plots reached anthesis (A) (VI_A) and 2) postanthesis (PA) at the milk and dough developmental stages (VI_PA).

**Table 1.** Spectral vegetation indices assessed in this study.

| Vegetation index | Band centre (nm) | Spectral band | References |
| --- | --- | --- | --- |
| *Structural indices* |  |  |  |
| Normalized Difference Vegetation Index (NDVI) | 790, 660 | NIR, Red | (Rouse et al. 1974) |
| Renormalized Difference Vegetation Index (RDVI) | 790, 660 | NIR, Red | (Roujean and Breon 1995) |
| Improved Soil Adjusted Vegetation Index (MSAVI) | 790, 660 | NIR, Red | (Qi et al. 1994) |
| Modified Triangular Vegetation Index (MTVI2) | 790, 660, 550 | NIR, Red, Green | (Haboudane et al. 2004) |
| Green Normalized Difference Vegetation Index (GNDVI) | 790, 550 | NIR, Green | (Gitelson et al. 1996) |
| *Chlorophyll indices* |  |  |  |
| Transformed CARI/Optimized Soil adjusted Vegetation Index (TCARI/OSAVI) | 790, 735, 660, 550 | NIR, Red Edge, Red, Green | (Haboudane et al. 2002) |

### Genotyping

The panel was genotyped with 13177 SNP markers using the Illumina Infinium 15K Wheat SNP Array at Trait Genetics GmbH (Gatersleben, Germany), and 11196 markers were ordered according to the SNP map developed by Wang et al. (2014). To reduce the risk of errors in further analyses, markers and accessions were analysed for the presence of duplicated patterns and missing values. After excluding markers with more than 25% missing values and with a minor allele frequency (MAF) lower than 5%, a total of 10090 SNPs were used for mapping purposes.

### Statistical analyses

Restricted maximum likelihood (REML) was used to estimate the variance components and to produce the best linear unbiased predictors (BLUPs) for agronomic traits and VIs, following the MIXED procedure of the SAS-STAT statistical package (SAS Institute Inc., Cary, NC, USA). To assess differences between years and genetic subpopulations, one-way ANOVAs were conducted for the whole collection. Least squares means were calculated and compared using the Tukey HSD test at *p*<0.05. Mean phenotypic values across the three years were used to perform a hierarchical cluster analysis by the Ward method (Ward 1963). All analyses were carried out using the JMP v14.2.0 statistical package (SAS Institute, Inc., Cary, NC, USA), considering a significance level of *p*<0.05.

### Marker trait associations

A GWAS with 10090 SNP markers was conducted on the whole germplasm collection using Tassel 5.0 software (Bradbury et al. 2007) for all agronomic and VI traits per year and across the three growing seasons. A mixed linear model (MLM) was fitted using a principal component analysis (PCA) matrix with 6 principal components as the fixed effect and a kinship (k) matrix as the random effect (PCA + K model) at the optimum compression level. In addition, the anthesis date was incorporated as a cofactor in the analysis, as reported in previous studies (Crowell et al. 2016; Condorelli et al. 2018; Soriano et al. 2021). Manhattan plots were generated using the R script CMplot (https://github.com/YinLiLin/CMplot). A frequently used threshold was established at −log_10_ *P* > 3, as previously reported in the literature (Wang et al. 2014b, 2020; Mangini et al. 2018; Sukumaran et al. 2018; Condorelli et al. 2018). Confidence intervals (CIs) for MTAs were calculated for each chromosome according to the LD decay reported by Rufo et al. (2019). To simplify the MTA information, the associations were grouped into QTL hotspots. To define a hotspot, the density of MTAs along the chromosome was calculated as the QTL overview index (Chardon et al. 2004) for each cM.

### Gene annotation and *in silico* gene expression analysis

Gene annotation for the target region of QTL hotspots was performed using the gene models for high-confidence genes reported for the wheat genome sequence (IWGSC 2018), available at https://wheat-urgi.versailles.inra.fr/Seq-Repository/Annotations. Physical distances were estimated using the genetic distances from the markers flanking the CIs of each QTL hotspot.

*In silico* expression analysis and the identification of upregulated gene models were carried out using the RNA-seq data available at http://www.wheat-expression.com/ (Ramírez-González et al. 2018) for the following studies: 1) drought and heat stress time-course in seedlings, 2) spikes with water stress, 3) seedlings with PEG to simulate drought, and 4) shoots after two weeks of cold.

Gene Ontology (GO) data were retrieved from the high-confidence gene annotation at https://wheat-urgi.versailles.inra.fr/Seq-Repository/Annotations.

## Results

### Environmental conditions

The experimental site has a typical Mediterranean climate characterized by an irregular pattern of yearly rainfall distribution, low temperatures in winter that rise sharply in spring and high temperatures continuing until the end of the crop cycle. Figure 1 represents a graphical summary of the rainfall and maximum and minimum temperatures during the crop cycle across the three years of field trials. All the weather variables were representative of long-term data from the region for each growing season, although 2017 was considered exceptionally dry due to the low rainfall received. The last year (2018) was characterized as the wettest from December (sowing) to June (physiological maturity) with 269 mm of rainfall, whereas the first and second growing seasons with 207 mm and 105 mm of rainfall were rather dry, respectively, suffering severe water scarcity during the grain filling period with only 5 mm of precipitation.

**Figure 1.**
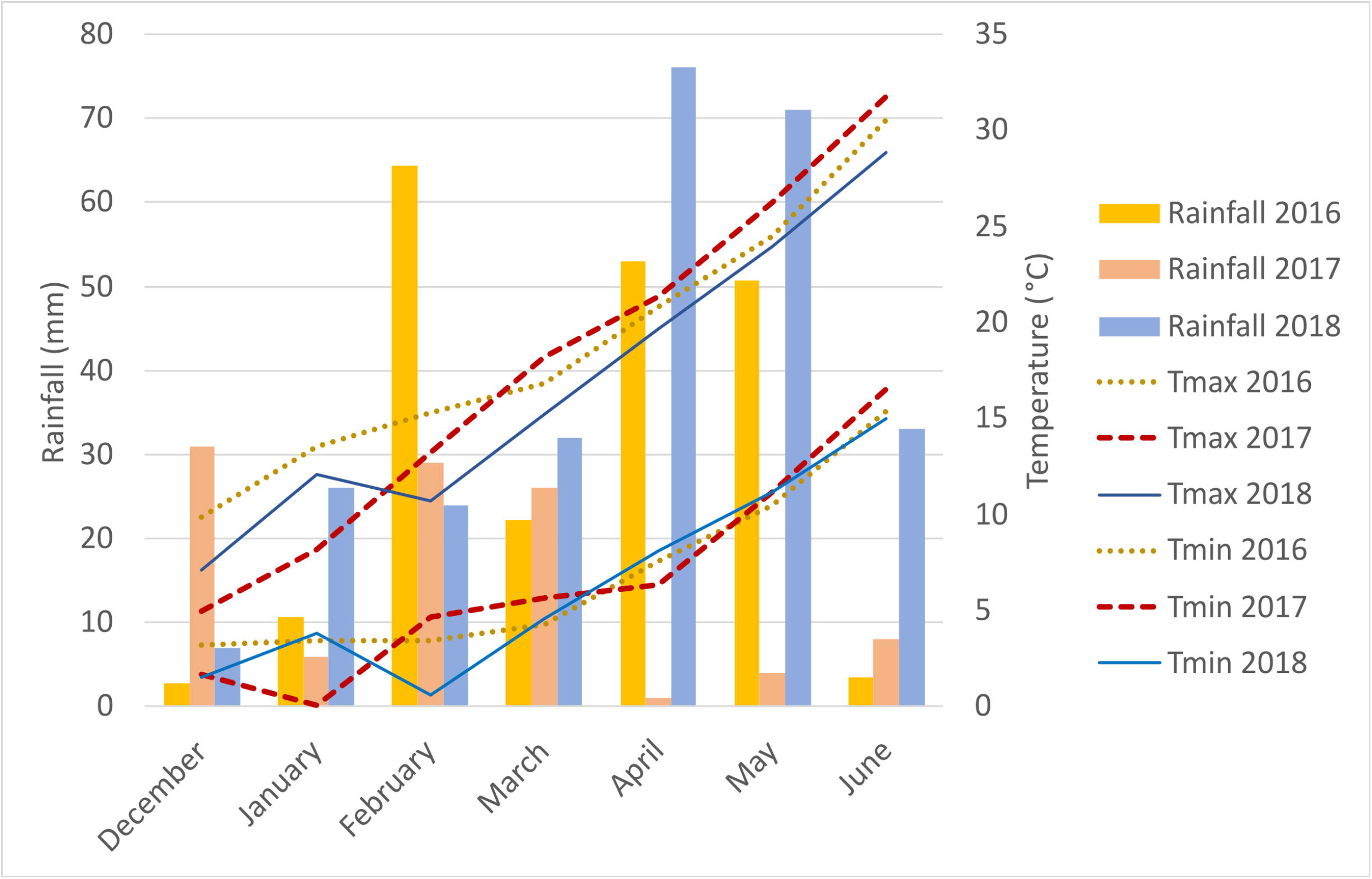
Monthly rainfall (mm) and minimum (Tmin) and maximum (Tmax) temperatures during the growth cycle of each growing season.

### Phenotypic analyses

The results of the analyses of variance (ANOVAs) for the agronomic traits measured during the three growing seasons are shown in Table 2. The percentage of variability explained by year was the highest for GA (81.6%) and GS65 (67.9%), while the sum of squares of SP was the highest for yield (76.5%), NGm^2^ (65.0%), HI (62.6%), PH (61.4%) and GS87 (59.0%). Finally, the highest percentage explained by the interaction between year and SP was found for biomass, NSm^2^ and TKW, reaching 86.2%, 84.9% and 71.0%, respectively. Significant differences were found between SPs and years for all traits. The year x SP interaction was also significant for all traits, except for HI.

**Table 2.** Analysis of variance for grain yield, harvest index (HI), biomass, number of spikes per square metre (NSm^2^), number of grains per square metre (NGm^2^), thousand kernel weight (TKW), plant height (PH), green area (GA), number of days from sowing to anthesis (GS65), and grain filling duration (GFD, GS87) for the three years of field trials. SS, sum of squares; SP, subpopulation. * *p* < 0.01. ** *p* < 0.001.

| SS (%) | Yield | HI | Biomass | NSm<sup>2</sup> | NGm<sup>2</sup> | TKW | PH | GA | GS65 | GS87 |
| --- | --- | --- | --- | --- | --- | --- | --- | --- | --- | --- |
| Year | 1.3** | 0.1 | 5.2** | 8.0** | 6.5** | 10.5** | 2.6** | 81.6** | 67.9** | 8.7** |
| SP | 76.5** | 62.6** | 8.6** | 7.1** | 65.0** | 18.5** | 61.4** | 2.2** | 30.4** | 59.0** |
| Year x SP | 22.2** | 37.3 | 86.2** | 84.9* | 28.5** | 71.0* | 36.0** | 16.2** | 1.7** | 32.3** |

Table 3 shows the results of the ANOVA for the VIs and LAI estimated through the MTVI2 at the anthesis and PA stages during 2017 and 2018. Differences between subpopulations and between years, as well as the year x SP interaction, were statistically significant for all traits in both years. The sum of squares for year accounted for 1.3% (NDVI) to 92.9% (RDVI) of the variation at anthesis, whereas at PA, the percentages ranged from 10.0% (TCARI/OSAVI) to 92.3% (LAI). The percentages of the total variation explained by SP ranged from 2.3% (RDVI) to 11.0% (TCARI/OSAVI) at anthesis, while they ranged from 1.0% (MTVI2) to 11.2% (TCARI/OSAVI) PA. Year was the most important for explaining the variations in LAI, RDVI, MSAVI, MTVI2 and GNDVI in the two growth stages. SP explained the least percent of variation at both growth stages for all traits. The year x SP interaction accounted for 4.8% (RDVI) to 89.9% (NDVI) of the model variance at anthesis, with the highest values for NDVI and TCARI/OSAVI. The variance explained by the year x SP interaction at PA ranged from 6.6% (LAI) to 78.8% (TCARI/OSAVI).

**Table 3.** Analyses of variance for the LAI estimated through MTVI2 and all the VIs calculated at the anthesis stage and PA in 2017 and 2018. SS, % of the sum of squares; SP, subpopulation. * *p* < 0.01. ** *p* < 0.001.

| SS (%) | LAI | NDVI | RDVI | MSAVI | MTVI2 | TCARI/OSAVI | GNDVI |
| --- | --- | --- | --- | --- | --- | --- | --- |
| Anthesis |  |  |  |  |  |  |  |
| Year | 62.6** | 1.3* | 92.9** | 88.6** | 71.1** | 15.6** | 67.7** |
| SP | 8.9** | 8.8** | 2.3** | 2.7** | 8.9** | 11.0** | 8.0** |
| Year x SP | 28.5** | 89.9** | 4.8** | 8.7** | 20.0** | 73.4** | 24.3** |
| Post anthesis |  |  |  |  |  |  |  |
| Year | 92.3** | 85.3** | 88.2** | 90.8** | 91.8** | 10.0** | 83.4** |
| SP | 1.1** | 2.9** | 2.2** | 1.4** | 1.0** | 11.2** | 4.8** |
| Year x SP | 6.6** | 11.8** | 9.6** | 7.8** | 7.2** | 78.8** | 11.8** |

The mean values of phenotypic traits for each year and SP are shown in Table 4. Yearly means showed that the highest yield was in 2016, a year in which the yield components NSm^2^, NGm^2^ and TKW reached intermediate values between those obtained in the two subsequent years. The shortest duration of the preanthesis period and the longest GFD were also observed in 2016. On the other hand, the lowest yield, NSm^2^ and NGm^2^, the heaviest grains and the shortest GFD were observed in 2017. GA reached the highest value in 2016, which was characterized as the wettest year during the period from January-March, i.e., the stem elongation stage, when the trait was measured. In contrast, 2017 was the driest year in the same period, which showed the lowest value for GA. In 2017, PH showed maximum values but biomass showed the lowest values at maturity. Finally, in 2018, biomass, the number of spikes and grains per unit area showed high values, and the cycle until anthesis was the longest.

**Table 4.** Mean values of agronomic traits in a set of 170 landraces and 184 modern cultivars of bread wheat for each growing season and genetic subpopulation. Different letters at each growing season or subpopulation indicate significant differences at *p* ≤ 0.01 using Tukey’s honest significant difference test.

|  | **Yield**<br>**(t/ha)** | **HI** | **Biomass**<br>**(t/ha)** | **NSm<sup>2</sup>** | **NGm<sup>2</sup>** | **TKW**<br>**(g)** | **PH**<br>**(cm)** | **GS65** | **GA** | **GFD** |
| --- | --- | --- | --- | --- | --- | --- | --- | --- | --- | --- |
| 2016 | 8.0<sup>a</sup> | 0.35<sup>a</sup> | 17.5<sup>a</sup> | 532<sup>b</sup> | 20836<sup>a</sup> | 38.6<sup>b</sup> | 105.8<sup>b</sup> | 150<sup>c</sup> | 46.7<sup>a</sup> | 36<sup>a</sup> |
| 2017 | 7.4<sup>c</sup> | 0.36<sup>a</sup> | 16.1<sup>b</sup> | 509<sup>b</sup> | 17877<sup>b</sup> | 41.3<sup>a</sup> | 108.4<sup>a</sup> | 155<sup>b</sup> | 26.6<sup>c</sup> | 32<sup>c</sup> |
| 2018 | 7.9<sup>b</sup> | 0.36<sup>a</sup> | 18.1<sup>a</sup> | 583<sup>a</sup> | 21966<sup>a</sup> | 36.2<sup>c</sup> | 101.6<sup>c</sup> | 169<sup>a</sup> | 35.8<sup>b</sup> | 34<sup>b</sup> |
| SP1 | 5.2<sup>e</sup> | 0.29<sup>b</sup> | 16.6<sup>c</sup> | 556<sup>ab</sup> | 14065<sup>de</sup> | 38.2<sup>bc</sup> | 120.4<sup>a</sup> | 159<sup>b</sup> | 37.5<sup>a</sup> | 33<sup>c</sup> |
| SP2 | 5.9<sup>d</sup> | 0.30<sup>b</sup> | 16.7<sup>c</sup> | 534<sup>b</sup> | 15354<sup>d</sup> | 39.2<sup>b</sup> | 121.9<sup>a</sup> | 163<sup>a</sup> | 35.9<sup>bc</sup> | 31<sup>c</sup> |
| SP3 | 4.4<sup>f</sup> | 0.29<sup>b</sup> | 14.9<sup>d</sup> | 569<sup>ab</sup> | 13613<sup>e</sup> | 32.6<sup>d</sup> | 114.9<sup>b</sup> | 159<sup>b</sup> | 33.5<sup>d</sup> | 33<sup>c</sup> |
| SP4 | 10.8<sup>a</sup> | 0.44<sup>a</sup> | 18.5<sup>a</sup> | 568<sup>a</sup> | 28045<sup>a</sup> | 39.3<sup>b</sup> | 87.1<sup>d</sup> | 158<sup>b</sup> | 35.7<sup>c</sup> | 35<sup>b</sup> |
| SP5 | 9.2<sup>b</sup> | 0.43<sup>a</sup> | 18.0<sup>abc</sup> | 475<sup>c</sup> | 21872<sup>b</sup> | 43.2<sup>a</sup> | 92.1<sup>cd</sup> | 160<sup>b</sup> | 37.9<sup>a</sup> | 35<sup>b</sup> |
| SP6 | 9.7<sup>b</sup> | 0.42<sup>a</sup> | 18.0<sup>ab</sup> | 493<sup>c</sup> | 23877<sup>b</sup> | 42.0<sup>a</sup> | 95.7<sup>c</sup> | 152<sup>c</sup> | 38.1<sup>a</sup> | 37<sup>a</sup> |
| AD | 6.7<sup>c</sup> | 0.31<sup>b</sup> | 17.1<sup>bc</sup> | 550<sup>ab</sup> | 18414<sup>c</sup> | 36.9<sup>c</sup> | 111.4<sup>b</sup> | 157<sup>b</sup> | 36.8<sup>ab</sup> | 35<sup>b</sup> |

SP1: West Mediterranean landraces. SP2: North Mediterranean landraces. SP3: East Mediterranean landraces. SP4: Modern cultivars from France and Italy. SP5: Modern cultivars from Balkans. SP6: Modern cultivars from CIMMYT and ICARDA.

Significant differences in agronomic traits between subpopulations highlighted the division of the whole set into landraces and modern cultivars (Table 4). Modern SPs (SP4, SP5 and SP6) showed higher values of grain yield and yield components, HI, and biomass than landrace SPs. The highest value for grain yield was observed for SP4, in agreement with its higher number of spikes and grains per unit area. SP4 showed the lowest grain weight among modern SPs but was not significantly different from the heaviest grains observed in landraces (SP1 and SP2). As expected, landraces were taller than modern cultivars. SP3 showed the lowest value for GA. For phenology, SP2 took the longest time to reach the anthesis stage, whereas SP6 took the shortest time. In contrast, the GFD was the shortest for SP2 and the longest for SP6. Modern SPs showed a longer GFD than landraces.

The mean values of the VIs and LAI (estimated by MTVI2) at anthesis and at PA for 2017 and 2018 and the different SPs are shown in Tables 5 and 6, respectively. All traits had higher values at anthesis, except for TCARI/OSAVI. For all traits, differences between years were statistically significant at the two stages (Table 5). The LAI, RDVI and MSAVI showed the highest mean values in 2018. The mean values for TCARI/OSAVI were the highest in 2017. The year 2017 showed the highest values of MTVI2 and GNDVI at anthesis, but these VIs and NDVI were minimal at PA the same year. Due to saturation of the reflectance, NDVI became insensitive at high LAI values (LAI > 3) in both years at anthesis. LAI, NDVI, RDVI, MSAVI and MTVI2 significantly differed between landrace and modern cultivar SPs at anthesis, with higher values being recorded in the landraces (Table 6). However, no pattern was found for VI traits among SPs PA. SP2 and SP4 had higher mean values for all traits PA, with the exception of TCARI/OSAVI.

**Table 5.** Mean values of the LAI estimated by MTVI2 and all the VIs at anthesis and at PA in 2017 and 2018 from a set of 170 landraces and 184 modern bread wheat cultivars. Years followed by different letters indicate significant differences at *p* ≤ 0.01 using Tukey’s honest significant difference test.

|  | LAI | NDVI | RDVI | MSAVI | MTVI2 | TCARI/OSAVI | GNDVI |
| --- | --- | --- | --- | --- | --- | --- | --- |
| **Anthesis** |  |  |  |  |  |  |  |
| 2017 | 4.77<sup>b</sup> | 0.95<sup>a</sup> | 0.59<sup>b</sup> | 0.70<sup>b</sup> | 0.78<sup>a</sup> | 0.03<sup>a</sup> | 0.9<sup>a</sup> |
| 2018 | 6.29<sup>a</sup> | 0.94<sup>a</sup> | 0.78<sup>a</sup> | 0.91<sup>a</sup> | 0.98<sup>b</sup> | -0.51<sup>b</sup> | 0.84<sup>b</sup> |
| **Post anthesis** |  |  |  |  |  |  |  |
| 2017 | 1.56<sup>b</sup> | 0.65<sup>b</sup> | 0.42<sup>b</sup> | 0.43<sup>b</sup> | 0.39<sup>b</sup> | 0.07<sup>a</sup> | 0.62<sup>b</sup> |
| 2018 | 4.95<sup>a</sup> | 0.89<sup>a</sup> | 0.67<sup>a</sup> | 0.79<sup>a</sup> | 0.8<sup>a</sup> | -0.13<sup>b</sup> | 0.82<sup>a</sup> |

**Table 6.** Mean values of the LAI estimated by MTVI2 and all the VIs assessed at anthesis and in PA for each genetic subpopulation and the group of admixed genotypes from a set of 170 bread wheat landraces and 184 modern cultivars. Different letters at each subpopulation indicate significant differences at *p* ≤ 0.01 using Tukey’s honest significant difference test.

|  | LAI | NDVI | RDVI | MSAVI | MTVI2 | TCARI/OSAVI | GNDVI |
| --- | --- | --- | --- | --- | --- | --- | --- |
| **Anthesis** |  |  |  |  |  |  |  |
| SP1 | 5.88<sup>a</sup> | 0.96<sup>a</sup> | 0.71<sup>a</sup> | 0.83<sup>a</sup> | 0.92<sup>a</sup> | -0.08<sup>a</sup> | 0.87<sup>b</sup> |
| SP2 | 5.86<sup>a</sup> | 0.96<sup>a</sup> | 0.71<sup>a</sup> | 0.83<sup>ab</sup> | 0.92<sup>ab</sup> | -0.08<sup>a</sup> | 0.87<sup>b</sup> |
| SP3 | 5.75<sup>a</sup> | 0.95<sup>ab</sup> | 0.69<sup>b</sup> | 0.81<sup>c</sup> | 0.91<sup>ab</sup> | -0.02<sup>a</sup> | 0.87<sup>b</sup> |
| SP4 | 5.27<sup>b</sup> | 0.94<sup>bc</sup> | 0.67<sup>c</sup> | 0.78<sup>d</sup> | 0.85<sup>c</sup> | -0.63<sup>b</sup> | 0.89<sup>a</sup> |
| SP5 | 5.24<sup>b</sup> | 0.93<sup>c</sup> | 0.67<sup>c</sup> | 0.78<sup>d</sup> | 0.84<sup>c</sup> | -0.21<sup>a</sup> | 0.87<sup>bc</sup> |
| SP6 | 5.20<sup>b</sup> | 0.94<sup>c</sup> | 0.67<sup>c</sup> | 0.78<sup>d</sup> | 0.84<sup>c</sup> | -0.21<sup>a</sup> | 0.86<sup>c</sup> |
| AD | 5.70<sup>a</sup> | 0.95<sup>a</sup> | 0.69<sup>b</sup> | 0.81<sup>bc</sup> | 0.90<sup>b</sup> | -0.09<sup>a</sup> | 0.87<sup>b</sup> |
| **Post anthesis** |  |  |  |  |  |  |  |
| SP1 | 3.21<sup>bc</sup> | 0.76<sup>b</sup> | 0.56<sup>ab</sup> | 0.62<sup>ab</sup> | 0.61<sup>ab</sup> | 0.06<sup>ab</sup> | 0.70<sup>cd</sup> |
| SP2 | 3.32<sup>b</sup> | 0.80<sup>a</sup> | 0.57<sup>a</sup> | 0.64<sup>a</sup> | 0.64<sup>a</sup> | -0.08<sup>c</sup> | 0.73<sup>b</sup> |
| SP3 | 2.91<sup>d</sup> | 0.74<sup>cd</sup> | 0.54<sup>c</sup> | 0.60<sup>b</sup> | 0.59<sup>bc</sup> | 0.14<sup>a</sup> | 0.68<sup>d</sup> |
| SP4 | 3.50<sup>a</sup> | 0.79<sup>a</sup> | 0.55<sup>bc</sup> | 0.62<sup>b</sup> | 0.60<sup>b</sup> | -0.20<sup>d</sup> | 0.75<sup>a</sup> |
| SP5 | 3.36<sup>ab</sup> | 0.77<sup>bc</sup> | 0.53<sup>c</sup> | 0.60<sup>b</sup> | 0.59<sup>bc</sup> | -0.01<sup>abc</sup> | 0.73<sup>b</sup> |
| SP6 | 3.13<sup>c</sup> | 0.74<sup>d</sup> | 0.51<sup>d</sup> | 0.57<sup>c</sup> | 0.57<sup>c</sup> | 0.03<sup>abc</sup> | 0.70<sup>c</sup> |
| AD | 3.20<sup>bc</sup> | 0.76<sup>bc</sup> | 0.54<sup>c</sup> | 0.60<sup>b</sup> | 0.59<sup>b</sup> | -0.01<sup>bc</sup> | 0.71<sup>c</sup> |

To quantify the relation between trait variation and population structure, multiple linear regressions were carried out between population coefficients (Supplementary Table S2) and phenotypic performance for landrace and modern sets separately and both sets combined. The landrace R^2^ values ranged from 0.10 for MSAVI_A to 0.39 for GA, while the modern R^2^ values ranged from 0.10 for MTVI2_A to 0.64 for GNDVI_A. When the regressions were conducted on the combined data set, the R^2^ values ranged from 0.11 for biomass to 0.60 for NGm^2^. The traits yield and GNDVI_PA showed high R^2^ values (>0.35) for each set separately and for the combined set. The highest R^2^ values were found in modern set regressions for GNDVI_A, GNDVI_PA, NDVI_PA and GS65. Among the components of yield, TKW showed the highest R^2^ values in landrace set regressions, while in modern set regressions, NGm^2^ showed the highest R^2^ values.

The bidimensional clustering shown in Figure 2 represents the relationships among phenotypic traits and their mean values in the germplasm collection (3 years for agronomic traits and 2 years for VIs). The horizontal cluster grouped accessions according to their phenotypic similarity based on the traits in the vertical cluster. Horizontal clustering separated two main clusters: cluster A was composed only of landraces, and cluster B included modern cultivars and two landraces: *cv* ‘TRI 11548’ from Iraq and *cv* ‘1170’ from Turkey. Cluster A was characterized by lower yield and yield components, except NSm^2^, lower biomass, a shorter GFD but longer GS65, and taller plants than cluster B, but cluster A had higher values for VIs at anthesis, except for GNDVI_A.

**Figure 2.**
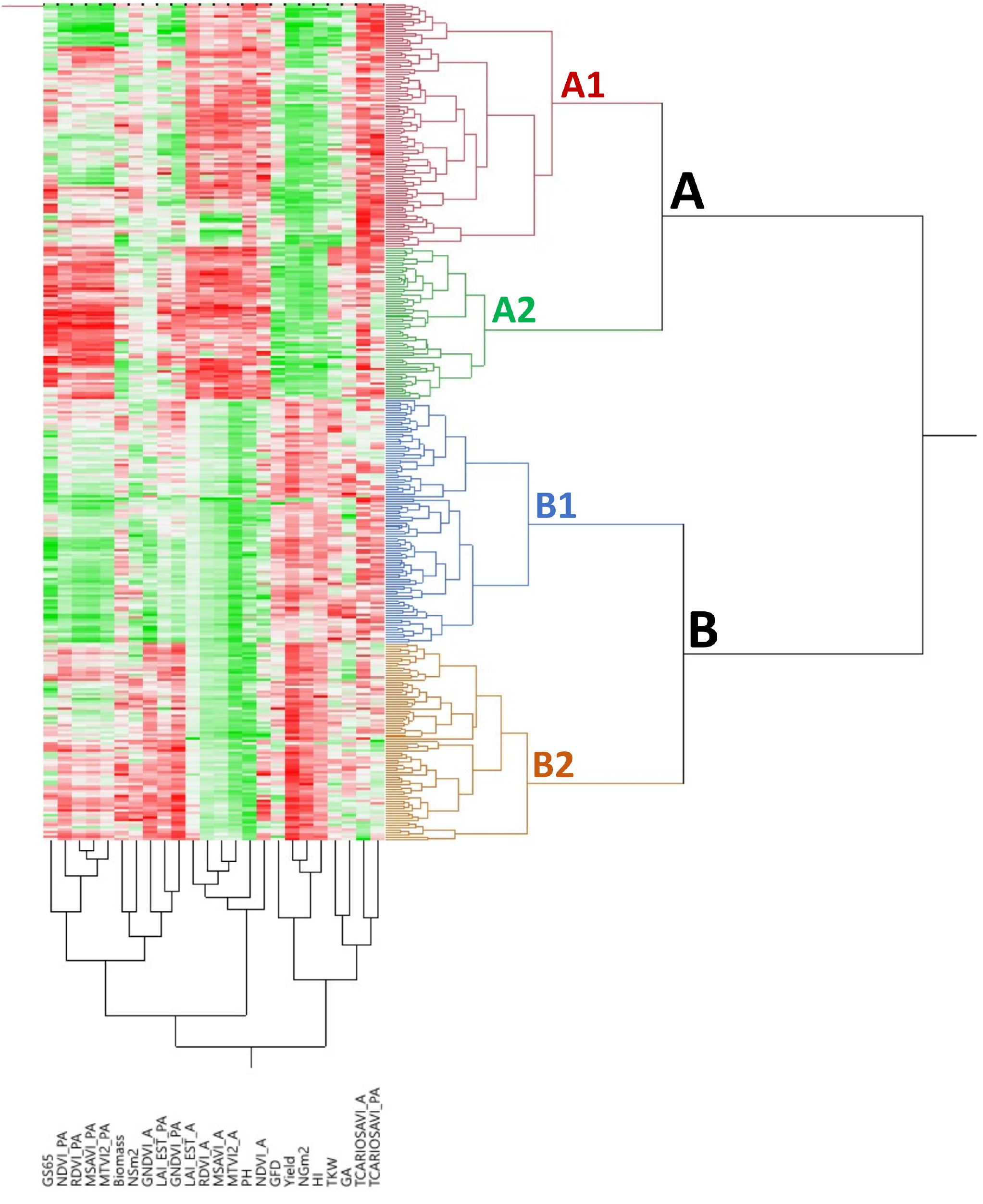
Bidimensional clustering showing the phenotypic relationships between the 354 bread wheat genotypes based on the analysed traits indicated in the vertical cluster at bottom. Red and green colours in the columns indicate high and low values, respectively. Dark: higher values; light: lower values; white: intermediate values.

Each of these two clusters was separated into two subclusters, A1 and A2 for landraces and B1 and B2 for modern cultivars. Subcluster A1 was represented mainly by south Mediterranean landraces (77%), including those from east and west regions, whereas subcluster A2 contained most of the north Mediterranean landraces (62%). East Mediterranean landraces were in a single cluster within A1, whereas west Mediterranean landraces were distributed in other clusters within A1. Differences among subclusters A1 and A2 were due to higher NSm^2^ and TCARIOSAVI_PA in A1 and longer cycles until anthesis in A2, along with higher values for GA and VIs at PA. Regarding modern cultivars, subcluster B1 was composed mainly of genotypes carrying the CIMMYT/ICARDA genetic background (SP6) (62%) and included the two landraces allocated to cluster B mentioned above. Moreover, subcluster B2 included 91% of the cultivars from SP4 (French and Italian modern cultivars). Most of the modern Balkan cultivars (SP5) were grouped in subcluster B1. Subcluster B1 was characterized by higher TCARIOSAVI_PA and GA, whereas subcluster B2 was characterized by higher NSm^2^, GNDVI_A and the rest of the VIs assessed in PA.

### Marker-trait associations

A summary of the results of the GWAS for all traits per year and for the mean values across years is reported in Figure 3. Using a common threshold of −log_10_ *P* > 3, as reported in the literature (Wang et al. 2017, 2020; Mangini et al. 2018; Sukumaran et al. 2018; Condorelli et al. 2018), a total of 2579 marker trait associations (MTA) were identified (Online resource 2). The year 2017 presented the highest number of MTAs, 74% of the total number of MTAs, whereas 2018 and the mean across years presented the lowest number of MTAs, 3% and 4%, respectively (Figure 3A). During 2016, only MTAs related to agronomic traits were reported, accounting for 19% of the total MTAs across years, as no multispectral images were captured during that year.

**Figure 3.**
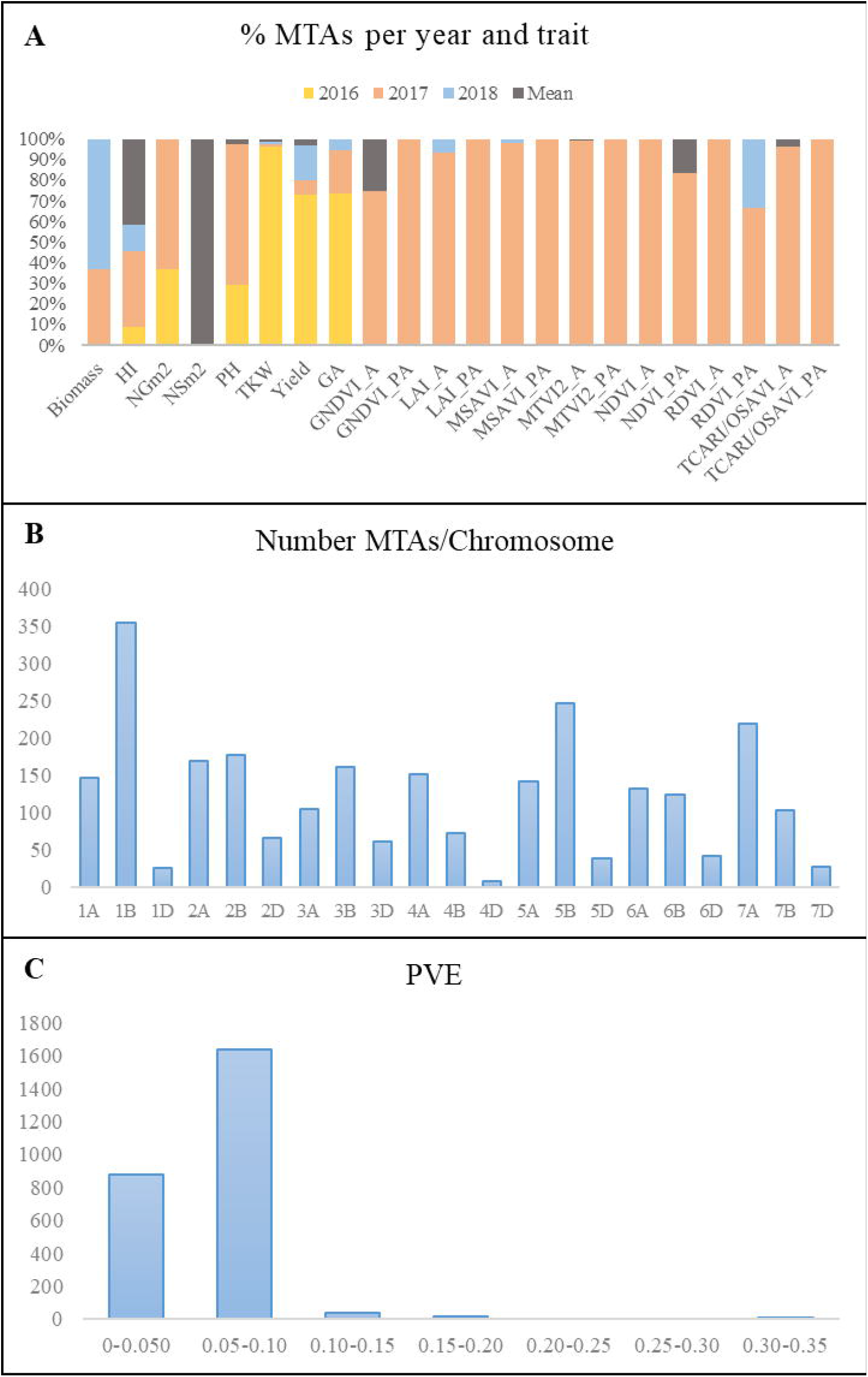
Summary of MTAs. (A) Percentage of MTAs per year and trait. (B) Number of MTAs per chromosome. (C) Phenotypic variance explained (PVE). HI, harvest index. LAI, leaf area index estimated by MTVI2. NSm^2^, number of spikes per square metre. NGm^2^, number of grains per square metre. TKW, thousand kernel weight (g). PH, plant height. GA, green area from emergence until the first node. A, anthesis stage. PA, post anthesis.

The number of MTAs per chromosome for all years and for the mean values across years ranged from 9 on chromosome 4D to 354 on chromosome 1B (Figure 3B). Genome B accounted for 48% of the total MTAs, followed by genomes A and D with 41% and 11%, respectively. The percentage of MTAs with a phenotypic variance explained (PVE) lower than 0.10 was 97.5%, which agreed with the highly quantitative nature of the analysed traits (Figure 3C).

A total of 815 MTAs were identified for seven agronomic traits (Table 8). Yield showed the highest number of MTAs (368), most of them (268) from 2016, whereas only one association was found for NSm^2^ with the mean across years. MTAs for TKW were found mainly during 2016 (96%), and those for PH were found mainly during 2017 (68%).

**Table 7.**
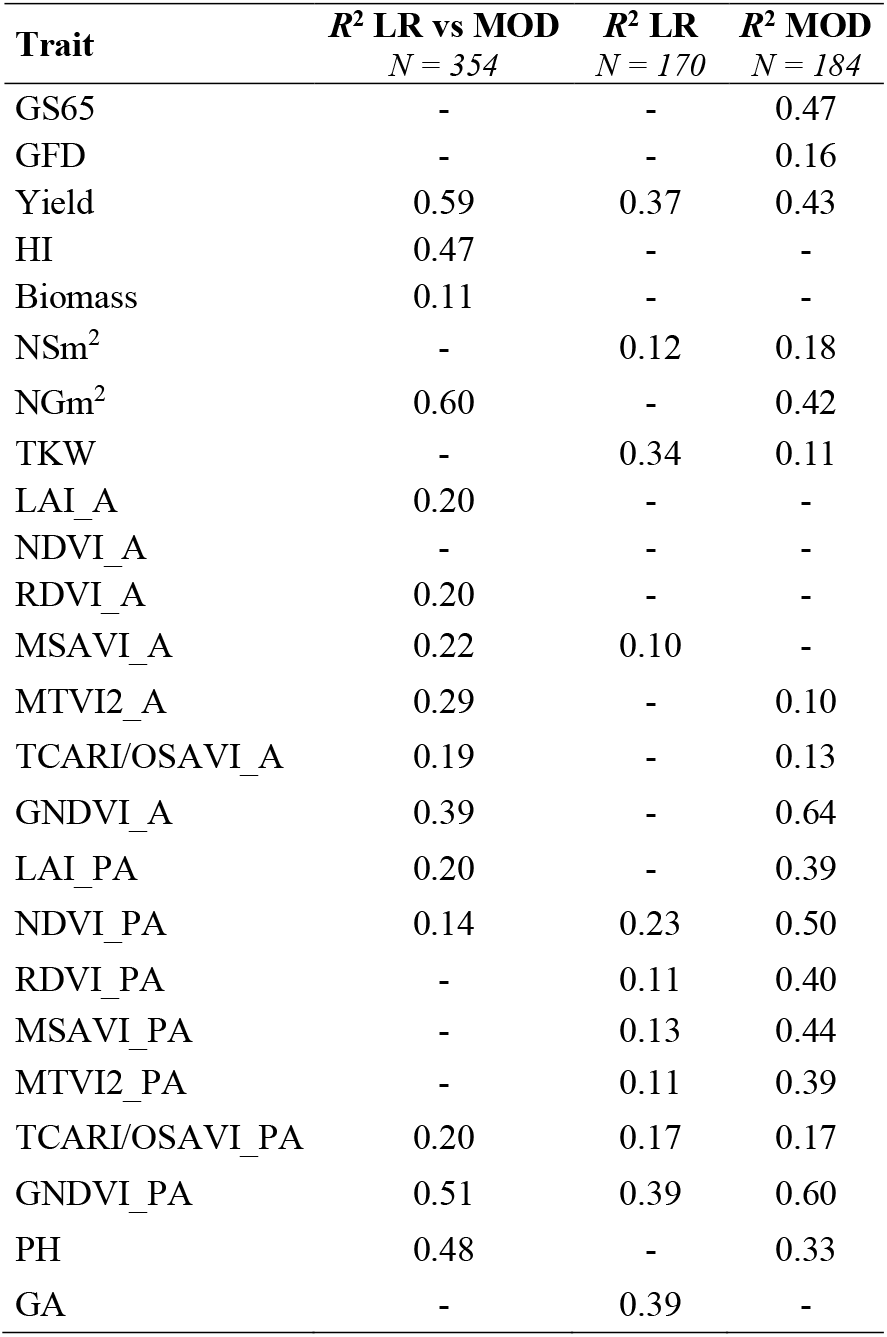
Relationship between trait variation and population structure (*q* values) for landrace and modern sets separately and the combined set.

**Table 8.** Total number of MTAs per year and trait with −log_10_*P* > 3

| Trait | 2016 | 2017 | 2018 | Mean | Total |
| --- | --- | --- | --- | --- | --- |
| Biomass | 0 | 7 | 12 | 0 | 19 |
| HI | 4 | 17 | 6 | 19 | 46 |
| NGm<sup>2</sup> | 11 | 19 | 0 | 0 | 30 |
| NSm<sup>2</sup> | 0 | 0 | 0 | 1 | 1 |
| PH | 65 | 150 | 0 | 6 | 221 |
| TKW | 125 | 2 | 1 | 2 | 130 |
| Yield | 268 | 26 | 62 | 12 | 368 |
| GA | 14 | 4 | 1 | 0 | 19 |
| GNDVI_A | - | 3 | 0 | 1 | 4 |
| GNDVI_PA | - | 8 | 0 | 0 | 8 |
| LAI_EST_A | - | 14 | 1 | 0 | 15 |
| LAI_EST_PA | - | 9 | 0 | 0 | 9 |
| MSAVI_A | - | 50 | 1 | 0 | 51 |
| MSAVI_PA | - | 3 | 0 | 0 | 3 |
| MTVI2_A | - | 348 | 0 | 2 | 350 |
| MTVI2_PA | - | 5 | 0 | 0 | 5 |
| NDVI_A | - | 1 | 0 | 0 | 1 |
| NDVI_PA | - | 5 | 0 | 1 | 6 |
| RDVI_A | - | 35 | 0 | 0 | 35 |
| RDVI_PA | - | 4 | 2 | 0 | 6 |
| TCARIOSAVI_A | - | 1195 | 1 | 47 | 1243 |
| TCARIOSAVI_PA | - | 9 | 0 | 0 | 9 |

A total of 1764 MTAs over −log_10_*P*>3 were identified for 15 VI traits (Table 8). Among them, 1718 were detected at or before anthesis (GA), and only 46 MTAs were identified PA. Ninety-six percent of the MTAs were identified during 2017, which was the year characterized by the lowest rainfall. TCARIOSAVI_A was the trait with the highest number of MTAs (1243), followed by MTVI2_A with 350.

To identify the genomic regions most involved in trait variation, QTL hotspots were identified using the QTL overview index defined by Chardon et al. (2004) for each cM of the genetic map. Confidence intervals were calculated using the LD decay for each chromosome reported by Rufo et al. (2019).

A total of 209 peaks were identified using the mean of the overview index across the 21 chromosomes (0.7) as the threshold, whereas using a high threshold (3.5), a total of 41 peaks were detected (Figure 4). These 41 peaks were reduced to 28 QTL hotspots (Online resource 3), 12 in genomes A and B and 4 in genome D. To simplify the search for candidate genes, QTL hotspots were excluded when 1) the centromere was included within the hotspot or the CI was higher than 35 Mb and 2) MTAs corresponded only to one year of field experiments. Eleven QTL hotspots grouping 295 MTAs remained for subsequent analysis (Table 9). As shown in Figure 4, hotspots defined by the QTL overview index correspond to genome regions with a higher number of MTAs.

**Figure 4.**
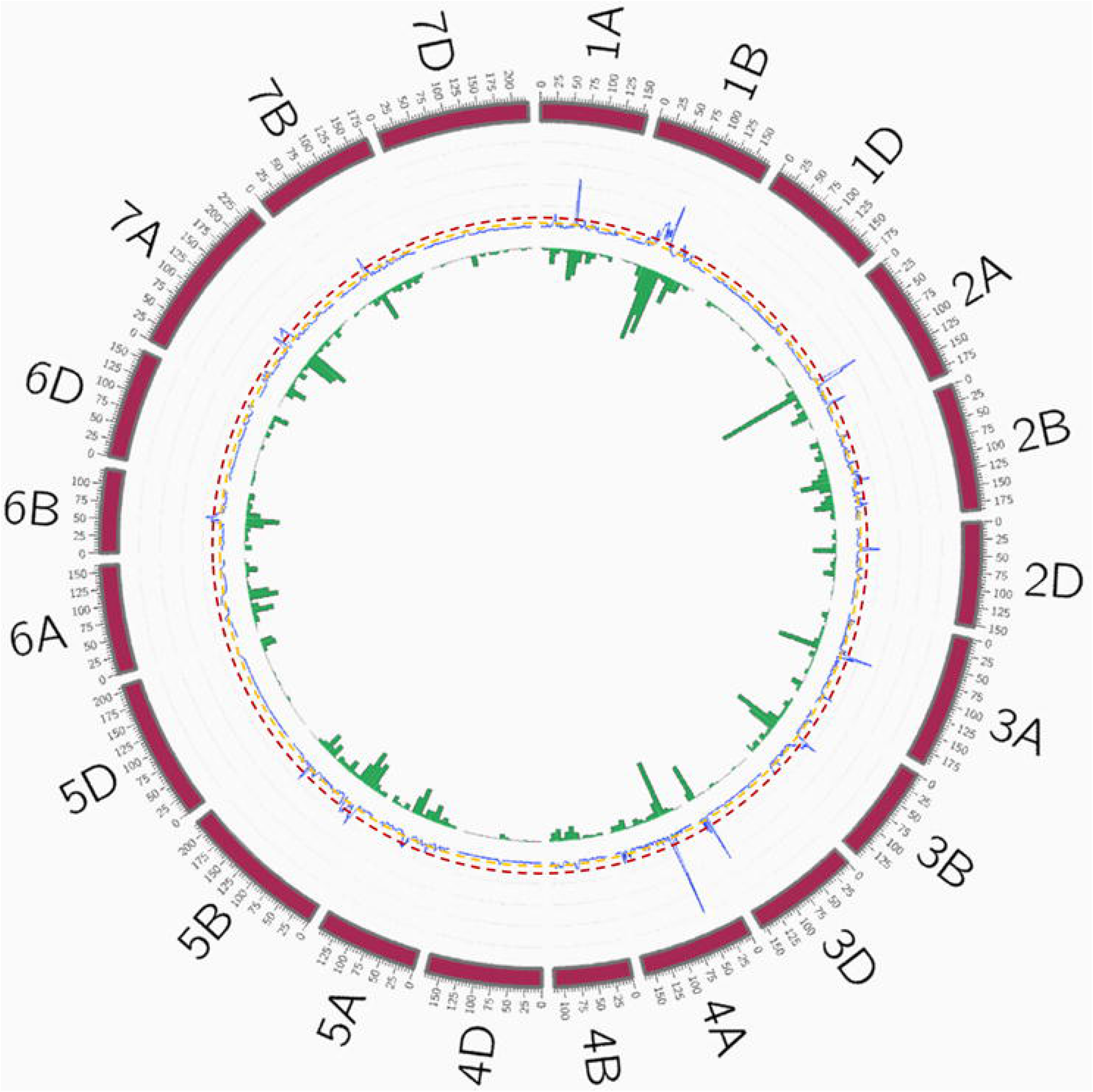
QTL overview index. The index values are represented along chromosomes as a blue line. Yellow and red dashed lines represent the thresholds for average (0.7) and higher values (3.5), respectively. Green bars below the QTL overview index represent the number of significant MTAs per 10 cM (−log_10_*P*>3).

**Figure 5.**
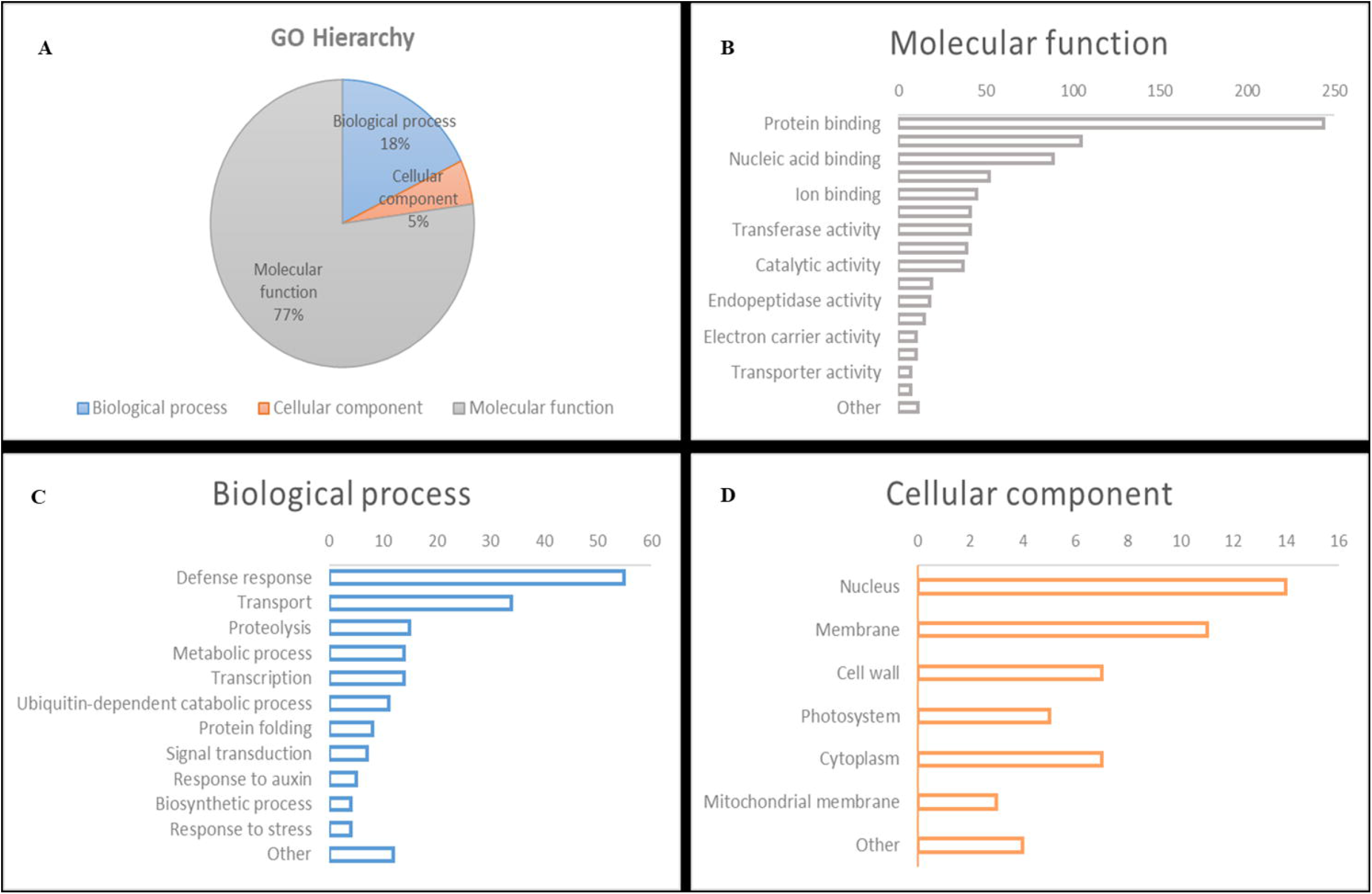
Gene Ontology classification of gene models within QTL hotspots.

**Table 9.** QTL hotspots identified for agronomic and remotely sensed VI-related traits. Positions are indicated in centimorgans (cM) and base pairs (bp). MTAs, marker trait associations; Env, number of environments; CI, confidence interval; A, anthesis; and PA, postanthesis.

| QTL Hotspot | Position (cM) | MTAs | Max -logP | Env | Number of Traits | Left marker | Position (bp) | Right marker | Position (bp) | CI (Mb) |
| --- | --- | --- | --- | --- | --- | --- | --- | --- | --- | --- |
| QTL1A.1 | 24-29-31 | 18 | 18.4 | 2 | 2 | BS00056550_51 | 7294564 | Kukri_c29655_239 | 9579957 | 2.3 |
| QTL1B.2 | 107-114-124 | 52 | 6.8 | 4 | 6 | BS00072791_51 | 629262942 | RFL_Contig2971_282 | 652455350 | 23.2 |
| QTL2A.2 | 149-151-153 | 25 | 5.5 | 2 | 6 | IAAV880 | 755788335 | Tdurum_contig50839_593 | 758514679 | 2.7 |
| QTL2D.1 | 39-41-45 | 29 | 6.2 | 2 | 6 | Kukri_c16477_181 | 61979485 | BS00067584_51 | 79416689 | 17.4 |
| QTL3A.2 | 176-177-178 | 9 | 5.9 | 2 | 5 | Excalibur_c77321_69 | 737299578 | Tdurum_contig31235_99 | 739397160 | 2.1 |
| QTL3D.1 | 142-143-144 | 32 | 7.0 | 2 | 5 | Kukri_rep_c87658_1436 | 613696030 | wsnp_Ex_c13629_21411429 | 609166802 | 4.5 |
| QTL4A.2 | 150-151-153 | 8 | 5.3 | 2 | 5 | Excalibur_c74390_108 | 733915685 | RAC875_c11702_1015 | 739524596 | 5.6 |
| QTL5B.2 | 54-61-64 | 51 | 6.2 | 3 | 6 | BobWHIte_c47103_84 | 457342562 | BS00039492_51 | 487602616 | 30.3 |
| QTL5B.3 | 68-69-69 | 10 | 5.4 | 3 | 3 | GENE-3574_643 | 519148851 | TA002629-0202 | 526576640 | 7.4 |
| QTL5B.4 | 159-161-163 | 21 | 18.2 | 3 | 8 | wsnp_Ku_c3151_5892200 | 680852782 | RAC875_c278_1801 | 683143220 | 2.3 |
| QTL7A.1 | 110-114-119 | 40 | 6.5 | 2 | 4 | BS00024786_51 | 79542753 | Kukri_rep_c101532_1046 | 84767559 | 5.2 |
| QTL Hotspot | Traits |  |  |  |  |  |  |  |  |  |
| QTL1A.1 | GNDVI_A, TCARI/OSAVI_A |  |  |  |  |  |  |  |  |  |
| QTL1B.2 | Yield, HI, MTVI2_A, PH, TCARI/OSAVI_A, TKW |  |  |  |  |  |  |  |  |  |
| QTL2A.2 | Yield, HI, LAI_A, MTVI2_A, TCARI/OSAVI_A, TKW |  |  |  |  |  |  |  |  |  |
| QTL2D.1 | Yield, MSAVI_A, MTVI2_A, PH, RDVI_A, TCARI/OSAVI_A |  |  |  |  |  |  |  |  |  |
| QTL3A.2 | Yield, LAI_PA, MTVI2_A, PH, TCARI/OSAVI_A |  |  |  |  |  |  |  |  |  |
| QTL3D.1 | Yield, LAI_PA, MTVI2_A, PH, TCARI/OSAVI_A |  |  |  |  |  |  |  |  |  |
| QTL4A.2 | Yield, MSAVI_A, PH, RDVI_A, TCARI/OSAVI_A |  |  |  |  |  |  |  |  |  |
| QTL5B.2 | Yield, HI, MSAVI_A, RDVI_A, TCARI/OSAVI_A, TKW |  |  |  |  |  |  |  |  |  |
| QTL5B.3 | Yield, HI, TCARI/OSAVI_A |  |  |  |  |  |  |  |  |  |
| QTL5B.4 | Yield, HI, LAI_A, MSAVI_A, MTVI2_A, PH, RDVI_A, TCARI/OSAVI_A |  |  |  |  |  |  |  |  |  |
| QTL7A.1 | Yield, MTVI2_A, PH, TCARI/OSAVI_A |  |  |  |  |  |  |  |  |  |

### *In silico* analysis of candidate genes

A search for CGs to study the relative gene expression levels under abiotic stress conditions and different tissues and developmental stages was performed within the QTL hotspot regions reported in Table 7 using the positions of flanking markers in the ‘Chinese Spring’ reference genome (IWGSC 2018) at https://wheat-urgi.versailles.inra.fr/Tools/JBrowse. A total of 1342 gene models were detected, and to classify this information, Gene Ontology (GO) for 1025 of the gene models (76%) was downloaded from https://wheat-urgi.versailles.inra.fr/Seq-Repository/Annotations (online resource 4). Seven hundred ninety-one CG were classified according to molecular function (MF), 183 according to biological process (BP) and 51 according to cellular component (CC). The most represented CGs according to molecular function were ‘protein binding’ (31%), ‘protein kinase activity’ (13%) and ‘nucleic acid binding’ (11%). According to BP, 30% of the CGs were involved in ‘defence response’ and 19% in ‘transport’. Finally, according to cellular component, 27% of the CGs were in the nucleus, 22% in the membrane and 14% in the cytoplasm and cell wall.

Subsequently, a search for differentially expressed genes (DEGs) upregulated under four abiotic stress conditions as reported in http://www.wheat-expression.com was carried out. These conditions included 1) drought and heat stress time-course in seedlings, 2) spikes with water stress, 3) seedlings treated with PEG to simulate drought, and 4) shoots after two weeks of cold, and DEGs were analysed in four tissues (roots, shoots/leaves, spikes and grains) during different developmental phases (seedling, vegetative and reproductive). A total of 12 CGs that were upregulated under abiotic stress were found in 6 QTL hotspots (Figure 6).

**Figure 6.**
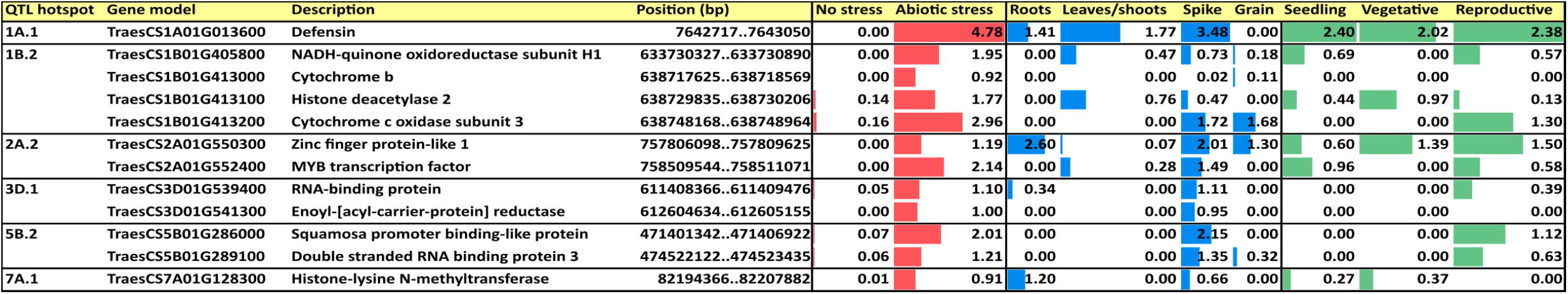
Upregulated candidate genes under abiotic stress conditions in 4 tissues and 3 developmental phases. Values are based on log_2_ (tpm) (transcripts per million).

Among the different DEGs, a defensin in hotspot QTL1A.1 showed the highest expression under abiotic stress conditions and was expressed in most of the tissues and all the developmental phases; it also showed the highest expression levels for each of the phases. All DEGs reported expression in the spikes with a range from 0.02 tpm for cytochrome b in QTL1B.2 to 3.48 tpm for defensin in QTL1A.1. Only four DEGs were expressed in the roots and five in the leaves/shoots and grain. Only zinc finger protein-like 1 in QTL2A.2 was expressed in all four plant tissues, showing the highest expression in roots. Regarding the developmental phase, no expression was reported in any stage for two DEGs, cytochrome b in QTL1B.2 and enoyl-[acyl-carrier-protein] reductase in QTL3D.1. The reproductive phase had the highest number of DEGs (9 out of 10 showing expression), whereas 6 were expressed in the seedlings and only 4 were expressed during the vegetative phase.

## Discussion

The current study was conducted under typical Mediterranean environmental conditions, with a pattern of increasing temperatures during the spring and an irregular distribution of rainfall across years. A GWAS panel of 354 bread wheat genotypes, including Mediterranean landraces and modern cultivars, was grown for three years under these conditions in northeastern Spain. Given that the decrease in the genetic diversity of wheat occurred during the second half of the 20^th^ century, associated with the introduction of high-yielding semidwarf cultivars (Autrique et al. 1996), landraces are considered a natural reservoir of genetic variation within the species and an invaluable source of new alleles to widen the genetic variability in breeding populations, particularly for traits regulating adaptation to suboptimal environments (Lopes et al. 2015). Recent studies have demonstrated the scarce use of wheat landraces in breeding programmes in the past, as suggested by the high genetic diversity and defined population structure among landrace and modern cultivar subpopulations (Soriano et al. 2016; Rufo et al. 2019).

### Phenotypic performance

The ANOVAs showed a large effect of SP on the phenotypic expression of the agronomic traits, whereas year showed the largest effect for most of the VIs, followed by the year x SP interaction, with the SP effect being the lowest. The variability in agronomic traits was mostly caused by the different agronomic performances of wheat landraces and modern cultivars, as reported in previous studies (Soriano et al. 2016; Royo et al. 2020). On the other hand, the high year effect on VIs was likely due to the contrasting water availabilities during the two years in which images were acquired by UAV in the experimental fields. This was not an unexpected result given that the decrease in the rate of growth of wheat caused by drought stress results in a severe reduction in total aboveground biomass (Royo et al. 2004; Villegas et al. 2014).

Yearly variation in weather conditions, particularly water input, resulted in a yield range from 7.4 t/ha in 2017, the driest year, to 7.9 and 8.0 t/ha during the years with higher rainfall. Even with the low water input, the average experimental yields were higher than expected in a severe drought environment. The numbers of spikes and grains per unit area were the highest in 2018, the wettest year, but were the lowest in 2017. Grain weight showed the opposite pattern, suggesting that under drier and hotter conditions, cultivars filled their grains at a higher rate (1.29 g/day in 2017 and 1.06 g/day in 2016 and 2018), thus showing a shorter GFD in 2017. The high yields recorded, considering the rainfed conditions of the field trials, could be attributed to the high soil fertility (approximately 3% of organic matter) and the superficial subsoil water layer at this site (Royo et al. 2021).

From a genetic viewpoint, a clear separation was observed between landraces and modern cultivars for most of the agronomic traits, which can be attributed to the improvement achieved by breeding. Among landraces, those from northern Mediterranean countries characterized by high rainfall and lower temperatures (Royo et al. 2014) showed higher yields due to an increase in the number of grains per unit area and grain weight. These genotypes showed longer cycles until anthesis and a shorter grain filling duration, although this last trait was not statistically significant. Landraces from eastern Mediterranean countries showed lower yields, a lower number of grains and lighter grains but an increase in the number of spikes per unit area compared with landraces from northern Mediterranean countries. Similar results for east Mediterranean landraces were previously reported in durum wheat by Soriano et al. (2018) and Roselló et al. (2019), suggesting an adaptation of landraces from this area to warmer environments, which has been associated with the allelic constitution of vernalization and photoperiod genes (Royo et al. 2020). The results of the current study are in agreement with previous research reporting a tendency for wheat to increase the number of ear-bearing tillers as an adaptation strategy under heat stress (Hütsch et al. 2019) and to increase the number of spikes per unit area in genotypes adapted to dry and warm areas compared with genotypes adapted to wetter and colder areas (Royo et al. 2014, 2020). Among modern cultivars, significant differences were mainly found between SP4 (cultivars from France and Italy) and the other two SPs (Balkans and CIMMYT-ICARDA-derived germplasm). These results suggest that breeding in France and Italy was in the direction of increasing yield through increasing the number of spikes and grains per unit area, whereas the other SPs showed higher TKW. In addition, the regression results of the modern set suggested a high impact of genetic population structure on the number of grains per unit area. Cultivars derived from CIMMYT and ICARDA germplasms reached anthesis earlier, up to eight days earlier compared with Balkan cultivars and six days earlier compared to French and Italian cultivars, which was in line with the high R^2^ values obtained in the relation between the modern set structure and GS65. This earliness can help these cultivars from warmer regions avoid heat stress at the end of flowering.

All traits related to HTP showed significant differences between years before and after anthesis, showing higher values for most of the VIs in 2018 than in the previous year. These higher values agree with the rainfall recorded for both years, which was significantly lower in 2017 than in 2018, mostly during the grain filling period. Furthermore, the difference in the mean values between growth stages was much higher in 2017. This result could be explained by the water scarcity particularly affecting the PA stage, which results in an important loss of chlorophyll content during the grain filling period; therefore, VIs using bands mostly placed in the near-infrared (NIR) and green regions showed lower values (Adamsen et al. 1999). Even though water stress affects the growth of wheat, the effects are higher during the grain filling period (Moragues et al. 2006). Thus, the LAI and GNDVI values decreased at the end of the growing cycle due to a low chlorophyll content associated with senescence during the grain filling period (Rufo et al. 2021). In addition, Gitelson et al. (2002) reported that the sensitivity of the green band was higher than that of the red band when the vegetation fraction was more than 60%, so vegetation indices using green wavelengths perform better at high LAI values, which in wheat under Mediterranean conditions are the highest at booting (Aparicio et al. 2000; Royo et al. 2004; Kyratzis et al. 2017; Rufo et al. 2021). TCARI/OSAVI were higher PA for both years. This agreed with the results of Zarco-Tejada et al. (2005), who reported that in advanced growth stages, chlorophyll indices such as TCARI performed better due to being less sensitive to the loss of turgor and leaf drop. In fact, these authors also stated that the different patterns of the indices across growth stages suggested that chlorophyll-related indices are more suitable closer to harvest, while structural indices related to canopy light scattering and growth are better for early stages.

Differences in the mean values of SPs were found in the two growth stages, with the highest values mainly found at anthesis based on the differences among years, thus highlighting the effect of PA senescence on the chlorophyll content. Landraces and modern cultivars showed significant differences in the LAI and structural VIs at anthesis, and these values were higher in the landraces. As reported in previous studies in durum wheat (García Del Moral et al. 2005; Soriano et al. 2018), landraces are characterized by their tolerance to water scarcity and their superior water use efficiency before anthesis compared to modern cultivars (Subira et al. 2015). Subpopulations showing the highest mean values for the LAI and VIs at PA were those including landraces from the north of the Mediterranean basin (SP2) and modern cultivars from France and Italy (SP4). Landraces from SP2 are better adapted to colder and wetter environments than landraces originating in the southern part of the Mediterranean basin. This adaptation pattern has been associated with the greatest early soil coverage and more aboveground biomass along the whole cycle length (Royo et al. 2014, 2021). For this reason, the canopy remains green much longer in landraces from northern Mediterranean countries than in those from southern Mediterranean countries (Royo et al. 2014). The same pattern was found in modern cultivars, with GNDVI values remaining higher than those of landraces after anthesis and being significantly different among modern subpopulations. These results agreed with those from the relationship between structure and GNDVI_PA, where the modern set showed the highest R^2^ values according to the differences found in GNDVI mean values among modern subpopulations. These results and the capacity to discern between landrace and modern SPs regarding the VI values at anthesis proved the accuracy of HTP in characterizing populations. Several studies have stated the potential of remote sensing for assessing agronomic traits by screening hundreds of plots in a short period of time, minimizing replications (Araus et al. 2018; Juliana et al. 2019; Gracia-Romero et al. 2019). Furthermore, various authors have stressed the suitability of using VIs measured early in the season for grain yield forecasting (Aparicio et al. 2000).

Bidimensional clustering was helpful to jointly visualize the results obtained by Tukey’s tests. Moreover, clustering of agronomic and HTP data revealed similarity with the separation obtained by Rufo et al. (2019) using SNP markers and SPs defined based on the structured collection. In both cases, a clear differentiation among landraces and modern cultivars was observed, which resulted in separation into two main clusters. Within the landrace cluster, (A) separation was observed between landraces from northern and southern Mediterranean countries, thus including landraces from SP2 in one cluster and those from SP1 and SP3 in the other cluster, with different groupings among them. Modern cultivars of SP6 (CIMMYT-ICARDA) clustered separately from the French and Italian cultivars (SP4), whereas modern cultivars from the Balkans grouped mostly with SP6. Although these two SPs were separated genetically, no significant differences were found for the agronomic traits, except for phenology, and regarding the VIs, no differences were found at anthesis. Two landraces (TRI 11548 and 1170) were included within modern cultivars from CIMMYT-ICARDA and the Balkans. These two landraces were characterized by a longer GFD, higher HI and lower number of spikes per unit area than the average for landraces. Landrace TRI 11548 from Iraq also showed higher yield and grain weight than other landraces, so it probably resulted from a selection made in an early landrace population.

### Marker trait associations

Dissecting the genetic basis of complex traits in plant breeding is essential to tackle molecular-based approaches for crop improvement. Several efforts have been previously made to identify QTLs and MTAs associated with traits of interest to carry out marker-assisted selection (MAS) approaches and the introgression of alleles of commercial interest in adapted phenotypes.

The highest number of MTAs related to agronomic traits was found in 2016, while 96% of MTAs related to VIs, GA and the LAI were identified in 2017. It has been reported that under contrasting conditions, the G×E interaction could affect the identification of stable associations among different environments (Mwadzingeni et al. 2017), which could explain the difference in the number of significant associations among the three years of field trials. The highest number of associations for yield and TKW in 2016 could be due to the moderate amount of water input (rainfall) during the spring, together with the longest grain filling duration, as reported in previous studies where grain weight predominantly enhanced yield in wet environments (García Del Moral et al. 2003; Moragues et al. 2006; Royo et al. 2006). Moreover, Royo et al. (2000) found that genotypes with longer GFDs could have greater opportunities to increase grain weight in favourable growing seasons than in warmer and drier seasons. The elevated number of VI-related MTAs found in the driest year (2017) could be explained by the higher variability in traits related to leaf biochemical properties or canopy structural attributes within the set of genotypes grown in environments with water scarcity (Rufo et al. 2021). The highest number of MTAs was identified for PH in 2017, when the tallest plants were observed. Qaseem et al. 2019 suggested that taller genotypes under drought stress could increase yield accumulation and convert more assimilates into grain. Of the 1764 MTAs detected for VIs, 1718 were found at anthesis, with 1243 for TCARI/OSAVI. This result could be explained by the significant differences found between landraces and modern SPs at anthesis for traits related to HTP. The highest variability was found when comparing SP4 with the rest of the SPs for TCARI/OSAVI, which could explain the elevated number of MTAs for this trait. The distribution of the MTAs across genomes agreed with the results of Rufo et al. (2020), with a similar number in the A and B genomes (41 and 48%, respectively) and the remaining 11% in the D genome. These results are consistent with those of previous studies (Chao et al. 2010; Wang et al. 2014b; Gao et al. 2015), which attributed these values to the lower genetic diversity and higher LD found in the D genome of bread wheat compared with genomes A and B (Rufo et al. 2019).

### QTL hotspots

To reduce the complexity of the high number of identified MTAs, QTL hotspots were defined using the QTL overview index proposed by Chardon et al. (2004). Although this statistic was initially used for classical biparental QTL analysis, we adapted it to GWAS using the confidence intervals of the MTAs as the distance of LD decay for each of the chromosomes. As reported in Figure 4, QTL hotspots defined by the high-value threshold of the overview index corresponded to genome regions with a higher MTA density, thus supporting the suitability of this approach in GWAS. To identify genome regions previously mapped in locations similar to our QTL hotspots and to detect new loci controlling agronomic traits and VIs, a comparison with previous GWAS studies and/or meta-QTL analysis reporting yield- and VI-related traits was conducted. Seven of the eleven QTL hotspots have been described previously in the literature. When compared with the meta-QTL analysis reported by Liu et al. (2020) in bread wheat, the QTL hotspots QTL1B.2 and QTL2D.1 were located in similar positions as MQTL1B.7 and MQTL1B.8 and MQTL2D.3 and MQTL2D.4, respectively, controlling grain yield, grain number and TKW under drought and heat stress. QTL1B.2 was also in the homologous region of QTL IWB50693 in durum wheat controlling spike length (Anuarbek et al. 2020), QSN.caas-1BL controlling NSm^2^ identified by Gao et al. (2015) in bread wheat and IWB3330 controlling the normalized chlorophyll pigment ratio index (NCPI) identified by Gizaw et al. (2018) in bread wheat. Gao et al. (2015) also found three QTLs for TKW, chlorophyll content and NDVI located in a common region with the hotspot QTL5B.2. This QTL hotspot was also detected in a similar region as QTL yield/root_5B.1 controlling grain yield and shoot length identified by Rufo et al. (2020). QTL1B.2 and QTL5B.2 were found to have homology with several studies. This was an expected result, since they were the longest hotspots including the highest number of MTAs. The genomic regions for QTL hotspots QTL5B.4 and QTL7A.1 were also found in common with three QTLs identified by Anuarbek et al. (2020) controlling the number of fertile spikes and TKW in durum wheat under rainfed conditions. Hotspots QTL1A.1 and QTL2A.2 shared a common position with mtaq-1A.2 reported by Roselló et al. (2019a) in durum wheat and QTL yield/root_2A2 identified by Rufo et al. (2020) in bread wheat, respectively, controlling root-related traits and grain yield in bread wheat.

The detection of these regions in common with other studies opens the opportunity to pyramid different QTLs with pleiotropic effects in future breeding approaches. Moreover, the use of the reference genome sequence makes it possible to rapidly identify common molecular markers to be used in MAS.

### Candidate genes

Gene annotation from the ‘Chinese spring’ reference genome sequence (IWGSC 2018) allowed us to identify 1342 gene models within the eleven QTL hotspots. Candidate gene mining was performed by searching for DEGs upregulated under drought conditions in different tissues and developmental stages through *in silico* analysis at http://www.wheat-expression.com. Among the 11 QTL hotspots, five did not present DEGs under the selected conditions; thus, gene models reported there were not used for subsequent analyses.

In the remaining six QTL hotspots, five candidate genes were upregulated under drought stress. They have been previously reported in the literature to be involved in stress resistance. Among them, in QTL hotspot 1A.1, a defensin protein (TraesCS1A01G013600) was found to show the highest expression under drought stress. According to Kumar et al. (2019), although defensins are mainly involved in antifungal responses, the defensin gene *Ca-AFP* from chickpea in transgenic *Arabidopsis* plants was overexpressed under drought stress and induced a higher germination rate, root length and plant biomass. The gene model TraesCS1B01G413200 in QTL hotspot 1B.2 encodes a cytochrome c oxidase that, according to Budak et al. (2013), appears to be downregulated in wheat in drought stress environments, in contrast to the cytochrome c oxidase found in our hotspot. Two gene models enhancing drought and heat stress tolerance were found in QTL hotspot 2A.2: the gene model TraesCS2A01G550300 encoding a zinc finger protein, as reported by (Yoon et al. 2014) in poplar, and the gene model TraesCS2A01G552400 encoding a MYB transcription factor, which was described by Zhao et al. (2018). *TaMYB31* from wheat is transcriptionally induced by drought stress in transgenic *Arabidopsis* plants. Finally, in QTL hotspot 5B.2, a squamosa-binding protein was identified (TraesCS5B01G286000). These protein families have been found to be involved in several biological processes. Cao et al. (2019), in expression studies of the Squamosa binding protein from wheat *TaSPL16*, found that this gene was highly expressed in young panicles but expressed at low levels in seeds, in agreement with the expression profile of TraesCS5B01G286000 found in our study. The ectopic expression of *TaSPL16* in *Arabidopsis* produced a delay in the emergence of vegetative leaves and early flowering and affected yield-related traits. Other gene models upregulated under drought stress, as reported in the RNA-seq analysis from Ramírez-González et al. (2018), such as NADH-quinone oxidoreductase, cytochrome b, histone deacetylase 2, RNA-binding protein, enoyl-[acyl-carrier-protein] reductase, double stranded RNA binding protein 3 and histone-lysine N-methyltransferase, have not been related to drought stress tolerance in the literature, and further experiments are required to assess their expression under drought stress conditions.

## Supporting information

Online Resource 1

Online Resource 2

Online Resource 3

Online Resource 4

## Availability of data and material

All data supporting the findings of this study are available within the paper and within its supplementary materials published online.

## Code availability

Not applicable

## Authors’ contributions

Conceptualization, JMS; methodology, RR, AL, MSL, JB, JMS; formal analysis, RR, JB, JMS.; data curation, RR; AL, JMS; writing—original draft preparation, RR; writing—review and editing, RR, MSL, JB, JMS; supervision, JMS; project administration, MSL, JMS; funding acquisition, MSL, JMS. All authors have read and agreed to the published version of the manuscript.

## Key message

GWAS identified hotspots genome regions for agronomic and vegetation indices in wheat, and the in silico gene expression analysis identified potential candidate genes.

## Acknowledgements

Authors acknowledges Conxita Royo for its helpful revision of the manuscript and the contribution of the CERCA Program (Generalitat de Catalunya).

## Online Resources

ESM1: Genotypes used in this study.

ESM2: Marker trait associations.

ESM3: QTL hotspots.

ESM4: Gene models within QTL hotspots.

