## Supplementary material for "Identification of QTL hotspots affecting agronomic traits and high-throughput vegetation indices in rainfed wheat": Online Resource 1

ESM 1. List of accessions. SP: Subpopulation based on STRUCTURE analysis (Rufo et al. 2019).

| **Accession** | **Country** | **SP** | **Type** |
| --- | --- | --- | --- |
| TRI 1667 | Albania | SP2 | Landrace |
| TRI 2100 | Albania | SP2 | Landrace |
| TRI 1671 | Albania | SP2 | Landrace |
| TRI 1313 | Bulgaria | SP2 | Landrace |
| TRI 7819 | Bulgaria | SP2 | Landrace |
| TRI 7821 | Bulgaria | SP2 | Landrace |
| 408-IV/61 | Bosnia & Herzegovina | SP2 | Landrace |
| Moriborska | Bosnia & Herzegovina | SP2 | Landrace |
| Ranka | Bosnia & Herzegovina | SP2 | Landrace |
| TRI 10515 | Cyprus | Admixed | Landrace |
| TRI 10526 | Cyprus | SP3 | Landrace |
| TRI 10561 | Cyprus | Admixed | Landrace |
| TRI 10590 | Cyprus | SP1 | Landrace |
| TRI 10531 | Cyprus | SP3 | Landrace |
| Ali Ben Makhloul | Algeria | SP3 | Landrace |
| Khalof | Algeria | SP3 | Landrace |
| Mahon Demias | Algeria | SP1 | Landrace |
| MG 17956 | Algeria | SP3 | Landrace |
| MG 17999 | Algeria | SP2 | Landrace |
| MG 18006 | Algeria | SP2 | Landrace |
| MG 18013 | Algeria | SP1 | Landrace |
| MG 18036 | Algeria | SP1 | Landrace |
| MG 18049 | Algeria | SP1 | Landrace |
| Bahatane | Algeria | SP3 | Landrace |
| Krelof - A | Algeria | SP2 | Landrace |
| Ble' du dahra-baal | Algeria | SP1 | Landrace |
| Mahon 7295 | Algeria | SP1 | Landrace |
| Sachah | Egypt | SP2 | Landrace |
| Gibson | Egypt | SP3 | Landrace |
| Hauch | Egypt | SP3 | Landrace |
| Hindiffino crible | Egypt | Admixed | Landrace |
| Hindiffino non crible | Egypt | SP3 | Landrace |
| Mokhtar | Egypt | SP1 | Landrace |
| Hindi 62 | Egypt | SP3 | Landrace |
| Bladette de Besplas | France | SP2 | Landrace |
| Ble Blanc de la Reole | France | SP2 | Landrace |
| Mars Rouge Sans Barbe | France | SP2 | Landrace |
| Touzelle Belle Abec | France | SP1 | Landrace |
| Touzelle Blanche Barbu | France | SP2 | Landrace |
| Touzelle Originario | France | Admixed | Landrace |
| Touzelle Rouge de Provence | France | Admixed | Landrace |
| Bladette de puylaurens | France | SP2 | Landrace |
| Mouton a epi rouge | France | SP2 | Landrace |
| Saisette | France | Admixed | Landrace |
| TRI 14046 | France | SP2 | Landrace |
| TRI 17938 | France | SP2 | Landrace |
| TRI 1425 | Greece | SP2 | Landrace |
| TRI 1686 | Greece | SP1 | Landrace |
| TRI 17989 | Greece | Admixed | Landrace |
| TRI 2060 | Greece | SP1 | Landrace |
| TRI 2071 | Greece | Admixed | Landrace |
| TRI 2129 | Greece | SP2 | Landrace |
| 986 | Croatia | SP2 | Landrace |
| Croatia 3 | Croatia | SP2 | Landrace |
| Croatia 6 | Croatia | SP2 | Landrace |
| Umarah | Iraq | SP1 | Landrace |
| TRI 15277 | Iraq | SP3 | Landrace |
| TRI 15292 | Iraq | SP3 | Landrace |
| TRI 16079 | Iraq | SP3 | Landrace |
| TRI 16084 | Iraq | SP2 | Landrace |
| TRI 11548 | Iraq | SP3 | Landrace |
| TRI 16080 | Iraq | SP3 | Landrace |
| TRI 11528 | Iraq | SP3 | Landrace |
| TRI 16063 | Iraq | SP3 | Landrace |
| TRI 8358 | Iraq | SP3 | Landrace |
| CGN06182 | Israel | SP1 | Landrace |
| CGN06204 | Israel | SP1 | Landrace |
| CGN04191 | Israel | Admixed | Landrace |
| Palestinskaya | Israel | Admixed | Landrace |
| 17310 | Israel | SP1 | Landrace |
| Cappellina | Italy | SP2 | Landrace |
| TRI 15321 | Italy | SP1 | Landrace |
| TRI 14055 | Italy | SP2 | Landrace |
| TRI 15226 | Italy | Admixed | Landrace |
| TRI 16900 | Italy | SP2 | Landrace |
| TRI 16895 | Italy | SP3 | Landrace |
| TRI 14842 | Italy | SP1 | Landrace |
| Solina | Italy | SP2 | Landrace |
| TRI 15219 | Italy | Admixed | Landrace |
| TRI 14173 | Italy | SP2 | Landrace |
| TRI 16516 | Italy | SP2 | Landrace |
| Cappelli | Italy | SP1 | Landrace |
| 25 | Jordan | SP3 | Landrace |
| Dorziyeh Karak - B | Jordan | SP3 | Landrace |
| SY 271 | Jordan | Admixed | Landrace |
| 17411 | Jordan | SP1 | Landrace |
| Beyrouth 11 | Lebanon | SP3 | Landrace |
| Beyrouth 3 | Lebanon | SP2 | Landrace |
| Salamouni | Lebanon | SP3 | Landrace |
| TRI 17974 | Libya | Admixed | Landrace |
| TRI 14643 | Libya | SP3 | Landrace |
| TRI 14668 | Libya | SP3 | Landrace |
| Canivano | Morocco | SP1 | Landrace |
| Fez 2 | Morocco | SP1 | Landrace |
| Recio | Morocco | SP1 | Landrace |
| CGN04157 - A | Morocco | Admixed | Landrace |
| CGN04158 | Morocco | SP1 | Landrace |
| CGN06246 | Morocco | SP1 | Landrace |
| CGN06247 | Morocco | SP1 | Landrace |
| CGN06248 | Morocco | SP1 | Landrace |
| CGN06252 | Morocco | SP1 | Landrace |
| CGN06255 | Morocco | SP1 | Landrace |
| CGN06260 | Morocco | Admixed | Landrace |
| CGN06264 | Morocco | SP1 | Landrace |
| CGN06266 | Morocco | Admixed | Landrace |
| CGN06269 | Morocco | SP1 | Landrace |
| CGN06271 | Morocco | Admixed | Landrace |
| CGN06284 | Morocco | Admixed | Landrace |
| CGN06297 | Morocco | SP1 | Landrace |
| TRI 18287 | Morocco | SP1 | Landrace |
| TRI 18291 | Morocco | SP1 | Landrace |
| TRI 18308 | Morocco | SP1 | Landrace |
| 309-VII/33 | Makedonia | SP2 | Landrace |
| 340-VII/45 | Makedonia | SP2 | Landrace |
| VII/1-B | Makedonia | SP2 | Landrace |
| Stara bela | Makedonia | SP2 | Landrace |
| Magueija | Portugal | SP2 | Landrace |
| Santareno | Portugal | SP2 | Landrace |
| Temporao de coruche | Portugal | SP2 | Landrace |
| Tremes branco | Portugal | SP1 | Landrace |
| Bistra | Romania | SP2 | Landrace |
| Pades | Romania | SP2 | Landrace |
| Solonetu nou | Romania | SP2 | Landrace |
| SVGB10195 | Romania | SP2 | Landrace |
| Raton de Belalcazar | Spain | SP1 | Landrace |
| Cabezorro | Spain | SP1 | Landrace |
| Negrete de Cañaveras | Spain | SP2 | Landrace |
| Pelon blanco | Spain | Admixed | Landrace |
| Chamorro de Villadiego | Spain | SP2 | Landrace |
| Blat petit de Olot | Spain | SP1 | Landrace |
| Candeal | Spain | SP2 | Landrace |
| Isla de Fuerteventura | Spain | SP2 | Landrace |
| Hembrilla de Jerga | Spain | SP2 | Landrace |
| Extremo Sur Argelino | Spain | SP2 | Landrace |
| Xeixa Tarragona | Spain | SP1 | Landrace |
| 41-II/4-B | Serbia | SP2 | Landrace |
| Crvenica | Serbia | SP2 | Landrace |
| Piskulja | Serbia | SP2 | Landrace |
| Legan bez osja | Serbia | SP2 | Landrace |
| 401 | Syria | SP1 | Landrace |
| Aleppo 21 | Syria | Admixed | Landrace |
| Aleppo 28 | Syria | SP3 | Landrace |
| Aleppo 32 | Syria | SP3 | Landrace |
| Aleppo 33 | Syria | SP3 | Landrace |
| Damaskus 12 | Syria | SP3 | Landrace |
| Damaskus 8 | Syria | SP1 | Landrace |
| K1140 | Syria | Admixed | Landrace |
| Kaundouhari | Syria | SP3 | Landrace |
| TRI 8375 | Syria | SP1 | Landrace |
| Salamuni - A | Syria | Admixed | Landrace |
| Allorca | Tunisia | SP2 | Landrace |
| Sbei noir | Tunisia | SP1 | Landrace |
| TRI 17006 | Tunisia | SP3 | Landrace |
| TRI 17002 | Tunisia | Admixed | Landrace |
| Florence 193 | Tunisia | SP2 | Landrace |
| 763 | Turkey | Admixed | Landrace |
| 1170 | Turkey | Admixed | Landrace |
| 811 (B) | Turkey | SP3 | Landrace |
| 1552 | Turkey | SP2 | Landrace |
| 2103 | Turkey | SP3 | Landrace |
| 2933 | Turkey | SP3 | Landrace |
| 2936 | Turkey | SP3 | Landrace |
| Edirne | Turkey | SP3 | Landrace |
| Gemir - B | Turkey | SP3 | Landrace |
| Kirmizi kiluk - A | Turkey | SP3 | Landrace |
| Ormece | Turkey | SP3 | Landrace |
| Saribasak | Turkey | SP3 | Landrace |
| T-317 | Turkey | SP2 | Landrace |
| Yazlik | Turkey | SP3 | Landrace |
| Yumusak | Turkey | SP3 | Landrace |
| Takhar 96 | Afghanistan | SP6 | Modern cultivar |
| Ain abid | Algeria | Admixed | Modern cultivar |
| Adelaide | Canada | Admixed | Modern cultivar |
| Misir-2 | Egypt | SP6 | Modern cultivar |
| Misir-1 | Egypt | SP6 | Modern cultivar |
| Gemmeiza-10 | Egypt | SP6 | Modern cultivar |
| Sakha-69 | Egypt | SP6 | Modern cultivar |
| Sids-12 | Egypt | SP6 | Modern cultivar |
| Gemmeiza-11 | Egypt | SP6 | Modern cultivar |
| Sahel-1 | Egypt | SP6 | Modern cultivar |
| Sids 1 | Egypt | SP6 | Modern cultivar |
| Adagio | France | SP4 | Modern cultivar |
| Candelo | France | SP4 | Modern cultivar |
| Aviso | France | SP4 | Modern cultivar |
| Belsito | France | SP4 | Modern cultivar |
| Innov | France | SP4 | Modern cultivar |
| Andalou | France | SP4 | Modern cultivar |
| Fiorenzo | France | SP4 | Modern cultivar |
| Sensas | France | SP4 | Modern cultivar |
| Adhoc | France | SP4 | Modern cultivar |
| Charles peguy | France | SP4 | Modern cultivar |
| Trocadero | France | SP4 | Modern cultivar |
| Astral | France | SP4 | Modern cultivar |
| Aerobic | France | SP4 | Modern cultivar |
| Bramante | France | SP4 | Modern cultivar |
| Soissons | France | SP4 | Modern cultivar |
| Isengrain | France | SP4 | Modern cultivar |
| Soberbio | France | SP4 | Modern cultivar |
| Cipres | France | SP4 | Modern cultivar |
| Bologna | France | SP4 | Modern cultivar |
| Avelino | France | SP4 | Modern cultivar |
| Viriato | France | SP4 | Modern cultivar |
| Nogal | France | Admixed | Modern cultivar |
| Premio | France | SP4 | Modern cultivar |
| Diamento | France | SP4 | Modern cultivar |
| Rgt Somontano | France | SP4 | Modern cultivar |
| Lazaro | France | SP4 | Modern cultivar |
| Altamira | France | SP4 | Modern cultivar |
| Andino | France | SP4 | Modern cultivar |
| Arezzo | France | SP4 | Modern cultivar |
| Solehio | France | SP4 | Modern cultivar |
| Mecano | France | SP4 | Modern cultivar |
| Guadalete | France | SP6 | Modern cultivar |
| Bonpain | France | SP6 | Modern cultivar |
| Equilibre | France | SP4 | Modern cultivar |
| Soledad | France | SP4 | Modern cultivar |
| Tremie | France | SP4 | Modern cultivar |
| Bastide | France | SP4 | Modern cultivar |
| Aubusson | France | SP4 | Modern cultivar |
| Garcia | France | SP4 | Modern cultivar |
| Kumberri | France | SP4 | Modern cultivar |
| Akim | France | SP4 | Modern cultivar |
| Camargo | France | SP4 | Modern cultivar |
| Exotic | France | SP4 | Modern cultivar |
| CCB Ingenio | France | SP4 | Modern cultivar |
| Bueno | France | SP4 | Modern cultivar |
| Sublim | France | SP4 | Modern cultivar |
| Alhambra | France | SP4 | Modern cultivar |
| Bandera | France | SP4 | Modern cultivar |
| Inoui | France | SP4 | Modern cultivar |
| SY Moisson | France | SP4 | Modern cultivar |
| Raffy | France | SP4 | Modern cultivar |
| Aguila | France | SP4 | Modern cultivar |
| Rodrigo | France | SP4 | Modern cultivar |
| Sollario | France | SP4 | Modern cultivar |
| Alpino | France | SP4 | Modern cultivar |
| Galpino | France | SP4 | Modern cultivar |
| Carles | France | SP4 | Modern cultivar |
| Sorrial | France | SP4 | Modern cultivar |
| Royssac | France | SP4 | Modern cultivar |
| Rimbaud | France | SP4 | Modern cultivar |
| Rvalo | France | SP4 | Modern cultivar |
| Sobbel | France | SP4 | Modern cultivar |
| Sofru | France | SP4 | Modern cultivar |
| Apache | France | SP4 | Modern cultivar |
| Illico | France | SP4 | Modern cultivar |
| Galopin | France | SP4 | Modern cultivar |
| Sobred | France | SP4 | Modern cultivar |
| Sokal | France | SP4 | Modern cultivar |
| Cezanne | France | SP4 | Modern cultivar |
| Craklin | France | SP4 | Modern cultivar |
| Paledor | France | SP4 | Modern cultivar |
| Botticelli | France | SP4 | Modern cultivar |
| Eureka | France | SP4 | Modern cultivar |
| MV Emese | Hungary | SP5 | Modern cultivar |
| Masaccio | Italy | SP4 | Modern cultivar |
| Zanzibar | Italy | SP4 | Modern cultivar |
| Palesio | Italy | SP4 | Modern cultivar |
| Toskani | Italy | SP4 | Modern cultivar |
| Trofeo | Italy | Admixed | Modern cultivar |
| Anapo | Italy | SP6 | Modern cultivar |
| Andana | Italy | SP4 | Modern cultivar |
| Anforeta | Italy | SP6 | Modern cultivar |
| Arabia | Italy | SP4 | Modern cultivar |
| Carisma | Italy | Admixed | Modern cultivar |
| Agape | Italy | SP4 | Modern cultivar |
| Antille | Italy | SP4 | Modern cultivar |
| Tiepolo | Italy | SP4 | Modern cultivar |
| Abate | Italy | SP4 | Modern cultivar |
| Arz | Lebanon | SP6 | Modern cultivar |
| Olga | Makedonia | SP5 | Modern cultivar |
| Balkania | Makedonia | SP5 | Modern cultivar |
| Siete cerros | Mexico | SP6 | Modern cultivar |
| Marchouch 8 | Morocco | SP6 | Modern cultivar |
| Aguilal | Morocco | SP6 | Modern cultivar |
| Achtar | Morocco | SP6 | Modern cultivar |
| Nesma | Morocco | SP6 | Modern cultivar |
| Arrehane | Morocco | SP6 | Modern cultivar |
| KG100 | Serbia | SP5 | Modern cultivar |
| PKB Arena | Serbia | SP5 | Modern cultivar |
| Ana Morava | Serbia | SP5 | Modern cultivar |
| PKB Lepoklasa | Serbia | SP5 | Modern cultivar |
| PKB Ratarica | Serbia | SP5 | Modern cultivar |
| Zvezdana | Serbia | SP5 | Modern cultivar |
| PKB Vizeljka | Serbia | SP5 | Modern cultivar |
| Simonida | Serbia | SP5 | Modern cultivar |
| BG Merkur | Serbia | SP5 | Modern cultivar |
| BG Carica | Serbia | SP5 | Modern cultivar |
| PLB Talas | Serbia | SP5 | Modern cultivar |
| BG Vitka | Serbia | SP5 | Modern cultivar |
| Vizija | Serbia | SP5 | Modern cultivar |
| PKB Mlinarka | Serbia | SP5 | Modern cultivar |
| Zlatna | Serbia | SP5 | Modern cultivar |
| Aleksandra | Serbia | SP5 | Modern cultivar |
| Planeta | Serbia | SP5 | Modern cultivar |
| Pobeda | Serbia | SP5 | Modern cultivar |
| Aurelia | Serbia | SP5 | Modern cultivar |
| Zemunska rosa | Serbia | SP5 | Modern cultivar |
| Renesansa | Serbia | SP5 | Modern cultivar |
| Kruna | Serbia | Admixed | Modern cultivar |
| NS 40S | Serbia | SP4 | Modern cultivar |
| Chambo | Spain | SP4 | Modern cultivar |
| 08THES2162 | Spain | SP6 | Modern cultivar |
| Vejer | Spain | SP6 | Modern cultivar |
| Antequera | Spain | SP6 | Modern cultivar |
| Conil | Spain | SP6 | Modern cultivar |
| Marchena | Spain | SP6 | Modern cultivar |
| Tejada | Spain | SP6 | Modern cultivar |
| Babui | Spain | SP6 | Modern cultivar |
| Catedral | Spain | SP6 | Modern cultivar |
| Eneas | Spain | SP6 | Modern cultivar |
| Califa sur | Spain | SP6 | Modern cultivar |
| Cartaya | Spain | SP6 | Modern cultivar |
| Escacena | Spain | SP6 | Modern cultivar |
| Jerezano | Spain | SP6 | Modern cultivar |
| Galeon | Spain | SP6 | Modern cultivar |
| Kilopondio | Spain | SP6 | Modern cultivar |
| Trebujena | Spain | SP6 | Modern cultivar |
| Victorino | Spain | SP6 | Modern cultivar |
| Alcala | Spain | SP6 | Modern cultivar |
| Rinconada | Spain | SP6 | Modern cultivar |
| Yecora | Spain | SP6 | Modern cultivar |
| Cielo | Spain | SP6 | Modern cultivar |
| Gazul | Spain | SP6 | Modern cultivar |
| Galera | Spain | SP6 | Modern cultivar |
| Mapeña | Spain | SP6 | Modern cultivar |
| Marca | Spain | SP6 | Modern cultivar |
| Anza | Spain | SP6 | Modern cultivar |
| Odiel | Spain | SP6 | Modern cultivar |
| Trimax | Spain | SP6 | Modern cultivar |
| Algido | Spain | Admixed | Modern cultivar |
| Artur Nick | Spain | SP6 | Modern cultivar |
| Mulhacen | Spain | SP6 | Modern cultivar |
| Platero | Spain | SP6 | Modern cultivar |
| Adalid | Spain | Admixed | Modern cultivar |
| Dollar | Spain | SP6 | Modern cultivar |
| Montcada | Spain | Admixed | Modern cultivar |
| Montserrat | Spain | Admixed | Modern cultivar |
| Idalgo | Spain | SP4 | Modern cultivar |
| Santoyo | Spain/France | SP4 | Modern cultivar |
| Debeira | Sudan | SP6 | Modern cultivar |
| Valbona | Switzerland | Admixed | Modern cultivar |
| Cham-8 | Syria | SP6 | Modern cultivar |
| Cham-6 | Syria | SP6 | Modern cultivar |
| Babaga-3 | Syria | SP6 | Modern cultivar |
| Cham-4 | Syria | SP6 | Modern cultivar |
| Hamam-4 | Syria | SP6 | Modern cultivar |
| Attila | Tunisia | SP6 | Modern cultivar |
| Karatopak | Turkey | SP6 | Modern cultivar |
| Ata 81 | Turkey | SP6 | Modern cultivar |
| Cumhuriyet 75 | Turkey | SP6 | Modern cultivar |
| Efe | Turkey | SP6 | Modern cultivar |
| Mane Nick | Turkey | SP6 | Modern cultivar |
| Gönen | Turkey | Admixed | Modern cultivar |
